# Electrostatic Actuation Induces Competing Adhesion and Vibration Regimes at Fingertip Contact

**DOI:** 10.64898/2026.01.06.697908

**Authors:** Celal Umut Kenanoglu, Michaël Wiertlewski, Yasemin Vardar

**Affiliations:** Department of Cognitive Robotics, Delft University of Technology, Delft, 2628 CD, The Netherlands

**Keywords:** electrostatic actuation, electrovibration, electroadhesion, contact mechanics, haptics

## Abstract

Electrostatic actuation enables programmable tactile feedback by modulating finger–surface friction via oscillating electric fields. Despite its potential, widespread adoption is hindered by an incomplete understanding of the underlying physical mechanisms, particularly the dynamics of finger–surface contact. To address this problem, this study presents the first time-resolved measurements of real contact area modulation under electrostatic actuation, obtained concurrently with contact forces. Experiments with ten participants sliding their fingers on an electrostatic display revealed an inverted U-shaped dependence of mean contact area and tangential force on actuation frequency, with a pronounced peak near 116 Hz—consistent with the frequency-dependent response of the fingertip–display system captured by mass–spring–damper and contact models. Two regimes emerged: a vibration regime below 320 Hz, where the voltage increased the contact area more than the tangential force, thereby reducing interfacial shear stress relative to the baseline; and an adhesion regime at higher frequencies, where skin viscoelasticity attenuated oscillations and restored or increased shear stress. For moist fingers, vibration effects were reduced, weakening the modulation of both tangential force and contact area. These findings reveal how adhesion and vibration jointly govern finger–surface interactions, guiding the design of next-generation electrostatic haptic interfaces.

---

If you have ever run your fingertip along the metal edge of a laptop, the chassis of a smartphone, or the base of a desk lamp connected to AC power, you may have felt a subtle, almost sticky change in friction. This peculiar sensation is caused by *electrostatic actuation*, an electrically induced attractive force at the skin–surface interface. First documented a century ago by Johnsen and Rahbek as *electroadhesion* under a constant DC voltage (*1*), this phenomenon was shown to increase friction between human skin and charged surfaces. Decades later, Mallinckrodt *et al*. (*2*) applied an alternating voltage to insulated metal electrodes and discovered that the resulting electrostatic force caused the finger to be periodically attracted to and released from the surface— an effect now known as *electrovibration*. Electrostatic actuation has since become a widely used technology for producing tactile feedback on surfaces, particularly on the touchscreens found in modern electronic devices (*3*).

Despite this long history, the mechanisms governing friction at the finger–surface interface under electrostatic actuation remain poorly understood. The increase in frictional force observed under such electrical loading is commonly attributed to an increase in real contact area, consistent with the adhesion model proposed by Bowden and Tabor (*4*). In this model, the kinetic friction is expressed as *F*_*t*_ = *τ A*, where *A* is the real contact area formed by microscopic asperity junctions, and *τ* denotes the interfacial shear stress during sliding (*5*). Prior studies have primarily interpreted the rise in friction by assuming that electrostatic loading increases *A*, while leaving *τ* constant (*6–8*), often relying on multiscale mean-field contact theory (*9, 10*) to infer area changes from measured tangential forces.

In haptic displays, however, electrostatic actuation is typically applied as an oscillating voltage at frequencies ranging from tens to hundreds of hertz to provide tactile feedback. The resulting electrostatic force therefore oscillates in time, periodically loading and unloading the fingertip and imposing a vibration on the contact. Although the magnitude of this motion—its effect on the finger contact area and resulting friction—inevitably depends on fingertip mechanics, most prior studies have primarily interpreted the frequency-dependent friction changes from an electrical perspective, attributing these changes to variations in the impedance of the finger–screen interface (*8, 11–13*). While this view explains how electrostatic pressure varies with frequency, it does not address how a fingertip responds to time-varying loading or how this loading alters instantaneous contact behavior during sliding. Recent measurements reveal a high-frequency attenuation in friction even when the electrical behavior is held approximately constant, consistent with a first-order, low-pass mechanical response of the fingertip (*14*). This attenuation points to a direct vibrational contribution to friction, in line with findings on fingertip contact under pure mechanical loading. For example, ultrasonic oscillations reduce sliding finger friction (*15*,*16*). Moreover, dynamic loading reduces the tangential force more than the contact area with increasing frequency, thus varying interfacial shear stress (*17*). Collectively, these findings indicate that vibration reshapes fingertip contact mechanics during sliding by attenuating friction that counteracts adhesion.

Taken together, these studies show that adhesion and vibration exert opposing effects on friction. Under oscillating electrostatic actuation on haptic touchscreens, these mechanisms operate simultaneously: the electric field generates an average attractive normal load that increases friction (adhesion), while its temporal modulation repeatedly pulls and releases the fingertip, imposing an oscillating motion on the contact (vibration). Their coexistence has led to a nearly interchangeable use of terms *(electro)adhesion*, referring to electrically induced attraction of the skin toward the surface and *(electro)vibration*, referring to oscillatory fingertip motion perceived by the user, even though they describe distinct physical phenomena. What remains unknown is how these antagonistic contributions, in combination with the fingertip’s mechanical response to the oscillating field, jointly determine the frequency dependence of contact area, tangential force, and interfacial shear stress under electrostatic actuation. Establishing this missing link is essential for explaining the mechanisms governing friction at the finger–surface interface and for designing haptic interfaces that control friction reliably through electrostatic fields.

Here, we show how these two fundamentally opposing phenomena, adhesion and vibration, jointly shape finger–surface contact under oscillating electric field by combining, for the first time, time-resolved friction measurements with synchronized imaging of contact area. The results indicate the existence of two distinct regimes, *vibration* and *adhesion*, with respect to frequency (Fig. 1). In the vibration regime, oscillatory motion enhances the contact area more than the tangential force, lowering interfacial shear stress (area-normalized tangential force) relative to the voltage-off reference. At higher frequencies, however, the increase in tangential force dominates, raising interfacial shear stress and marking a transition to an adhesion-dominated regime. This shift reflects the fingertip’s viscoelastic inability to follow rapid oscillations, a mechanism captured by a spring–damper model of the fingertip coupled with contact and friction models. Finally, all measured quantities exhibit substantial inter-participant variability, reflecting differences in fingertip mechanics and skin moisture.

**Figure 1:**
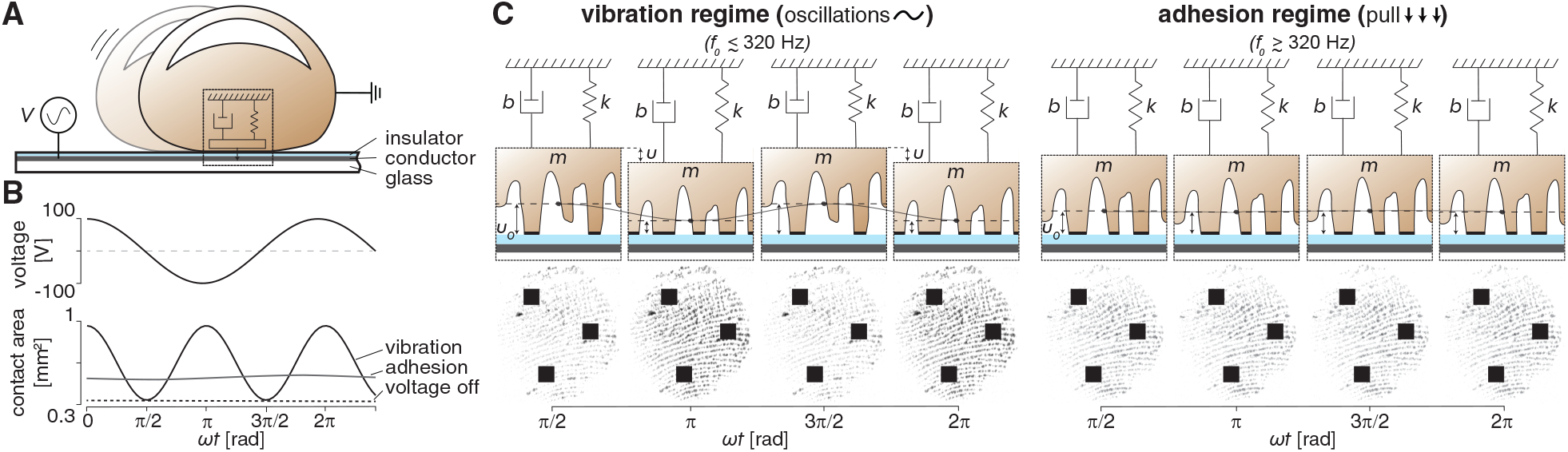
Frequency-dependent contact behavior under electrostatic actuation: vibration and adhesion regimes. (A) Schematic of a sliding fingertip on an electrostatic touchscreen, modeled as a lumped spring-mass-damper system. (B) Sinusoidal drive voltage and phase-dependent modulation of real contact area in two regimes; the dashed line indicates the 0-V baseline. (C) Response of the fingertip model in the normal direction, where *k, m*, and *b* represent spring, effective mass, and damping coefficients. Insets illustrate the micro-junction behavior in the two regimes: dynamic, oscillatory contact in the vibration regime and sustained, pulled-in contact in adhesion regime. The sinusoidal drive voltage *V* (*t*) = *V*_0_ cos(*ωt*), where *ω* = 2*π f*_0_ is the angular frequency corresponding to the input frequency *f*_0_ and *t* is time, is shown at its extrema and at a zero crossing. The dashed line in the insets marks the nominal interfacial separation, *u*_0_. At low frequencies ( *f*_0_ ≲ 320 Hz), vibration induces oscillatory skin motion, leading to phase-dependent modulation of real contact area. At higher frequencies, adhesion dominates, pulling the skin toward the surface and increasing contact area relative to the voltage-off reference.

## Results

We measured fingertip–touchscreen interaction in 10 participants under controlled conditions of normal force and speed, both with and without electrostatic actuation. Data were collected at ten logarithmically spaced frequencies (25–2500 Hz), with three repetitions per condition. An overview of the experimental setup and protocol, along with the recorded signals, is shown in Fig. 2; more details are provided in Materials and Methods and Figs. S1–S4.

**Figure 2:**
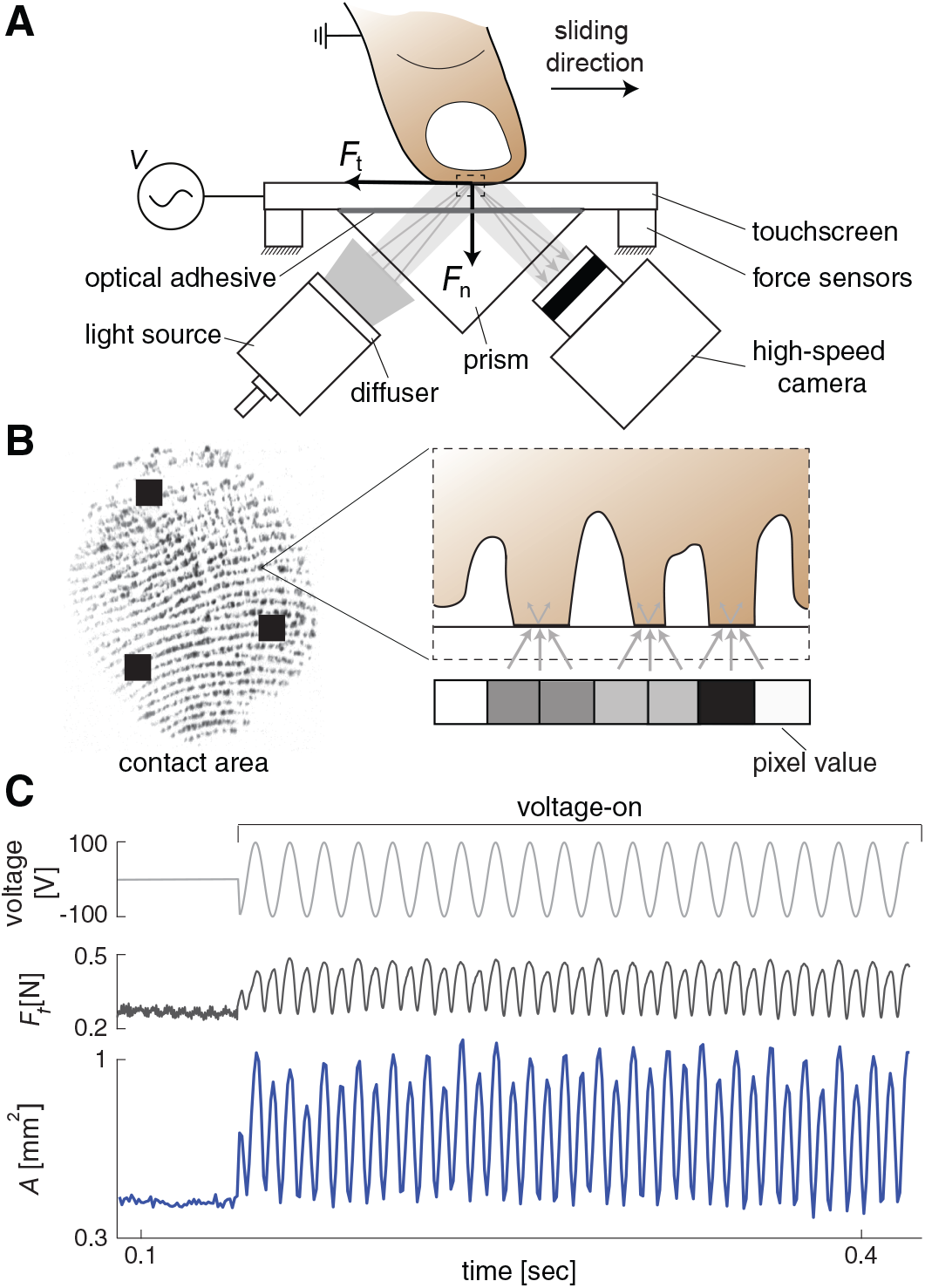
Experimental setup and representative measurements. (A) Schematic of the setup for simultaneous measurement of contact area and tangential force (*F*_*t*_) during finger sliding on a touchscreen under electrostatic actuation. A prism-based frustrated total internal reflection (FTIR) system illuminates the contact zone from below, allowing for direct imaging of fingertip contact using a high-speed camera and diffused light source. The touchscreen is optically bonded to the prism. Participants slid their finger at a constant normal force (*F*_*n*_ = 1 N) and speed (20 mm/s). During the initial one-third of each trial, the touchscreen was inactive (voltage-off), behaving as smooth glass, after which an alternating voltage (100 V peak) was applied to activate electrostatic actuation. Tangential force (*F*_*t*_), normal force (*F*_*n*_), fingertip contact area, and current were recorded throughout. (B) Principle of contact imaging with an example FTIR image. Light is totally internally reflected within the prism except at microscopic ridge asperities in intimate contact, where scattering causes dark pixels; non-contact regions remain bright. (C) Representative trial data showing input voltage, tangential force, and contact area over time.

Electrostatic actuation induced oscillations in both tangential force and real contact area at twice the input frequency *f*_0_, consistent with the quadratic dependence of electrostatic force on voltage (*13, 18*) (Figs. 2C and S5). Both signals reached maxima near the voltage extrema (±100 V) and minima at the zero-crossings (*V* = 0), where their values matched those measured under the voltage-off condition (Figs. 2C and 3A). Temporal modulation of real contact area over the drive cycle was clearly visible in the FTIR images, where darker regions correspond to greater contact (Fig. 3A; Movies S1 & S2).

**Figure 3:**
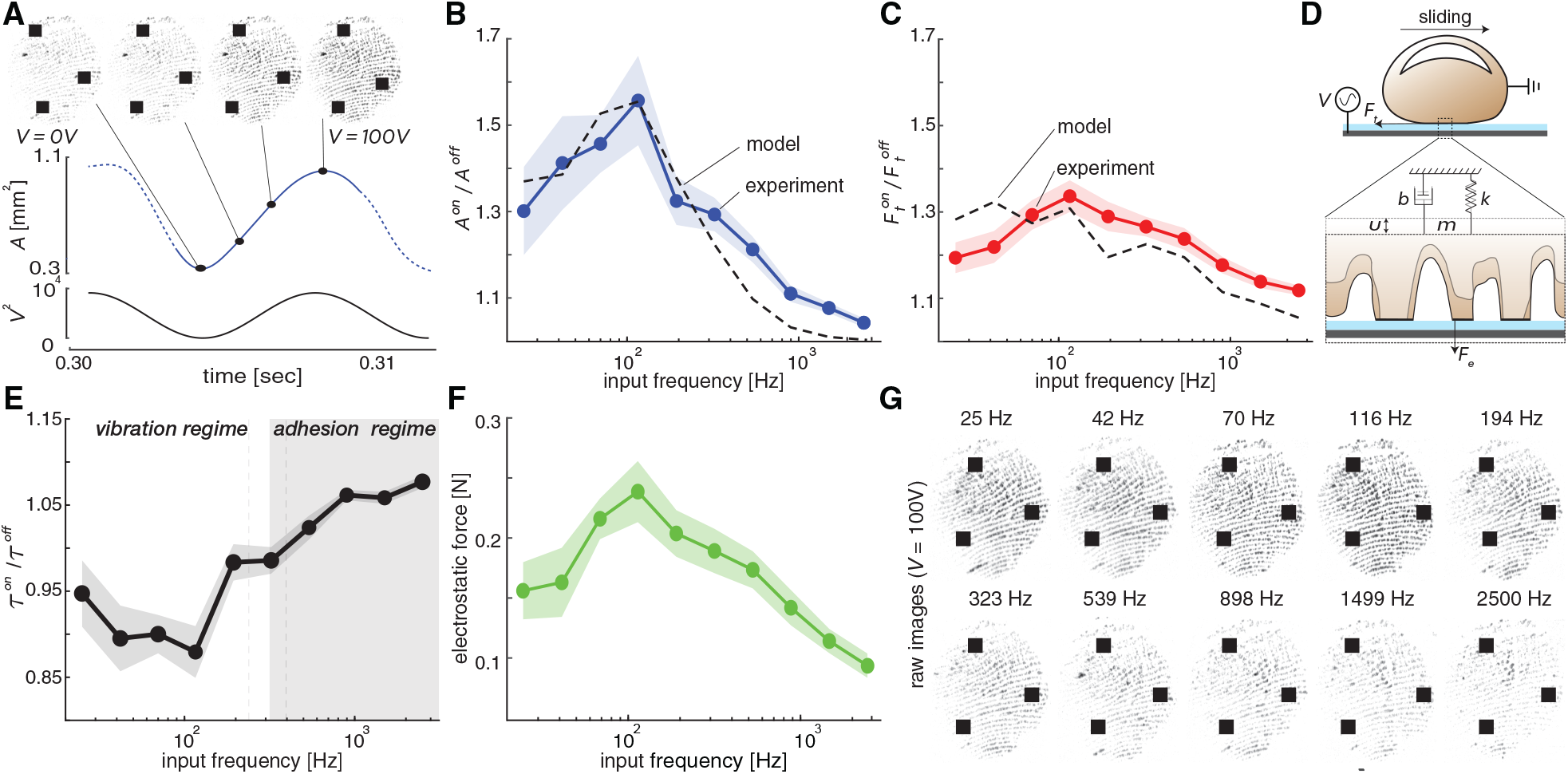
Experimental results. (A) Time evolution of the contact area at 70 Hz, illustrating oscillations induced by electrostatic actuation. (B-C) Ratio of measured contact areas (*A*^on^/*A*^off^) and tangential forces 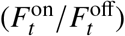 with and without electrostatic actuation across frequencies and corresponding model predictions (*9, 10, 15, 19, 20*). (D) Lumped spring–mass–damper model of the fingertip dynamics, where an electrostatic force *F*_*e*_ induces normal displacement *u* that increases the number of micro-junctions, thereby increasing the real contact area and tangential force, *F*_*t*_. (E) Interfacial shear stress ratio 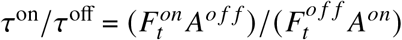. The transition zone around 320±82 Hz (between dashed lines) separates the vibration regime (*τ*^on^/*τ*^off^ < 1) from the adhesion regime (*τ*^on^/*τ*^off^ ≥ 1). (F) Estimated electrostatic force (1 − *μ*_off_/*μ*_on_)*F*_*n*_ (*8, 21, 22*). (G) Representative FTIR images of fingertip contact at different frequencies (100 V). Darker regions indicate greater contact, with stronger modulation in the vibration regime than in the adhesion regime. In B–F, large circles denote participant means, shaded bands indicate standard error of the mean, and superscripts “on/off” denote voltage on/off.

Changes in tangential force and real contact area across frequencies were quantified as the ratio of voltage-on to voltage-off values (i.e., with and without electrostatic actuation), computed within the same trial to ensure consistency. Throughout the manuscript, metrics averaged over the interval during which electrostatic actuation was applied are denoted by the superscript ‘on’, whereas those obtained without actuation are denoted by ‘off’. For both measures, these ratios exceeded unity at all tested frequencies, confirming the effect of electrostatic actuation (Figs. 3B & C and Figs. S6–S8). Moreover, they increased with frequency up to approximately 116 Hz, after which they declined. Below 320±82 Hz, the relative increase in contact area exceeded that of tangential force across participants, whereas above this frequency the trend reversed, with tangential force showing a slightly greater increase.

The measured real contact areas and tangential forces were compared with predictions from a spring–mass–damper model (*15*), a mean-field contact model based on Persson’s contact theory (*9, 10,15*), and a quasi-static model (*19*) to investigate whether these existing frameworks are consistent with the observed frequency-dependent behavior. The spring-mass-damper model was used to illustrate the fingertip’s mechanical response to electrostatic actuation in the normal direction, with the electrostatic force estimated as *F*_*e*_ = (1−*μ*_off_/*μ*_on_)*F*_*n*_, where *μ* is the measured friction coefficient and *F*_*n*_ is the applied normal load. The resulting vertical displacement, *u*, was then used as input to the mean-field and quasi-static models to estimate, respectively, the modulation of real contact area and tangential force. Then, the estimated contact area and tangential force ratios from the models were compared with the ones obtained from experimental measurements (Figs. 3B-D). In the mean-field contact model, the contact area depends on the microscale surface roughness and the applied normal pressure. The displacement *u* modulated the contact area as *A*^*on*^ = *A*^*o f f*^ exp(*u*_*m*_/*u*_*rms*_), where *u*_*rms*_ is the microscale root-mean-square roughness of the fingertip surface and *u*_*m*_ is the microscale displacement, obtained by scaling *u* with a dimensionless macro-to-micro factor. The resulting contact area exhibited a peak near 116 Hz and decreased at higher frequencies (Fig. 3B and Figs. S9–S11). The model output showed good agreement with the experimentally measured contact-area trend across frequencies, with relative errors ranging from 0.16% to 9.45% and a mean absolute error of 4.7% (Fig. 3B). The macroscale quasi-static model (*19*), which incorporates contact stiffness in normal and tangential directions, was used to examine the influence of normal vibration on tangential force. This model provides a physically interpretable approximation of the reduced increase in tangential force under oscillatory loading. Using the displacement estimated by the spring-mass-damper model, *u*, as input, the predicted tangential force trend closely matched the experimental trends, with errors ranging from 1.5% to 8.5% and a mean absolute error of 4.9% (Figs. 3C and Figs. S12-S13). However, the quasi-static model does not explicitly account for frequency-dependent viscoelastic and electrical effects; therefore, its validity is limited at higher frequencies, where these mechanisms become increasingly important. Further details of the model implementation are provided in the Supplementary Materials.

To assess the effects of frequency and participant variability, a linear mixed-effects model was employed for the *A*^on^/*A*^off^ and 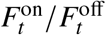 ratios. Frequency was modeled as a fixed effect and participant identity as a random intercept. The models revealed significant effects of frequency on both real contact area (*F* (9, 90) = 12.43, p < 0.001) and tangential force (*F* (9, 90) = 10.37, p < 0.001). Likelihood ratio tests comparing full models (with random effects) to reduced models (without) confirmed substantial inter-participant variability for both measures: real contact area (*χ*^2^(1) = 30.94, p < 0.001) and tangential force (*χ*^2^(1) = 23.43 , p < 0.001).

The pronounced frequency-dependent changes in real contact area and tangential force under electrostatic actuation indicated that interfacial shear stress was not constant, contradicting assumptions made in previous studies (*6–8*). The interfacial shear stress ratio, *τ*^*on*^/*τ*^*o f f*^ , calculated using the relation *F*_*t*_ = *τ A*, exhibited clear frequency-dependent variations (Figs. 3E and S14). Below 320±82 Hz, the increase in tangential force was less than the corresponding increase in contact area 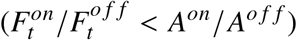, yielding *τ*^*on*^/*τ*^*o f f*^ < 1 and defining the *vibration* regime. This transition frequency represents the average across participants, with individual variability indicated by the light-gray shaded region in Fig. 3E. Above this frequency, *τ*^*on*^/*τ*^*o f f*^ ≥ 1, marking the onset of the *adhesion* regime. See Figs. S15–S18 for tangential force and contact area measurements from both regimes.

A linear mixed-effects model confirmed that the interfacial shear stress ratio (*τ*^*on*^/*τ*^*o f f*^ ) varied significantly with frequency (*F* (9, 90) = 19.49, p < 0.001). Post hoc paired t-tests with Benjamini-Hochberg correction revealed significant differences in *τ*^*on*^/*τ*^*o f f*^ between frequencies across the vibration and adhesion regimes (p<0.05), with the strongest contrast observed between the 70–116 Hz range and the adhesion regime (p<0.001). A likelihood-ratio test comparing a model including participant identity as a random effect to one without indicated a superior fit for the full model (*χ*^2^(1)=33.32, p< 0.001), confirming inter-participant variability. See Figs. S19–S21 for detailed pairwise results.

The electrostatic force also showed a frequency-dependent trend, rising to a peak near 116 Hz and then declining at higher frequencies (Fig. 3F). A linear mixed-effects model revealed a significant effect of frequency (*F* (9, 90) = 12.19, p < 0.001). A likelihood-ratio test comparing models with and without participant identity as a random effect confirmed significant inter-participant variability (*χ*^2^(1) = 44.99, p < 0.001). The shaded region in Fig. 3F illustrates this variability, which was more pronounced in the vibration regime.

Fingerprint images also revealed pronounced frequency-dependent variations in real contact area (see Fig. 3G for representative data from one participant). The intensity of the black regions, corresponding skin asperities at contact, was consistently greater at 100 V than at 0 V across all frequencies (see also Fig. 3A). The real contact area was larger in the vibration regime than in the adhesion regime, consistent with the stronger influence of electrostatic force at lower frequencies. To examine spatial patterns, we visualized fingerprint contact distributions with colormaps (Fig. S22), which show that electrostatic actuation primarily enhances central contact, which is already evident in the raw images (Figs. 3A & 3G). A qualitative comparison of the central and peripheral contact regions further indicated that the central enhancement was more pronounced in the vibration regime (Fig. S23). Consistent with this observation, a one-dimensional relative gap analysis along the middle of the fingerprint image showed that the difference between 100 V and 0 V remained relatively small near the contact edges but became more pronounced toward the center (Fig. S24). Representative videos of the fingertip contact during the vibration and adhesion regimes are provided in Movies S1–S4.

Lastly, we observed condensation in the fingerprint images of some participants (Fig. 4A and Movie S5), indicating the presence of finger moisture, as observed in previous studies (*23–26*). For these participants, both the changes in real contact area and tangential force between the voltage-on and voltage-off conditions were smaller (Fig. 4B). Correspondingly, the reduction in interfacial shear stress under electrostatic actuation was less pronounced compared to participants

**Figure 4:**
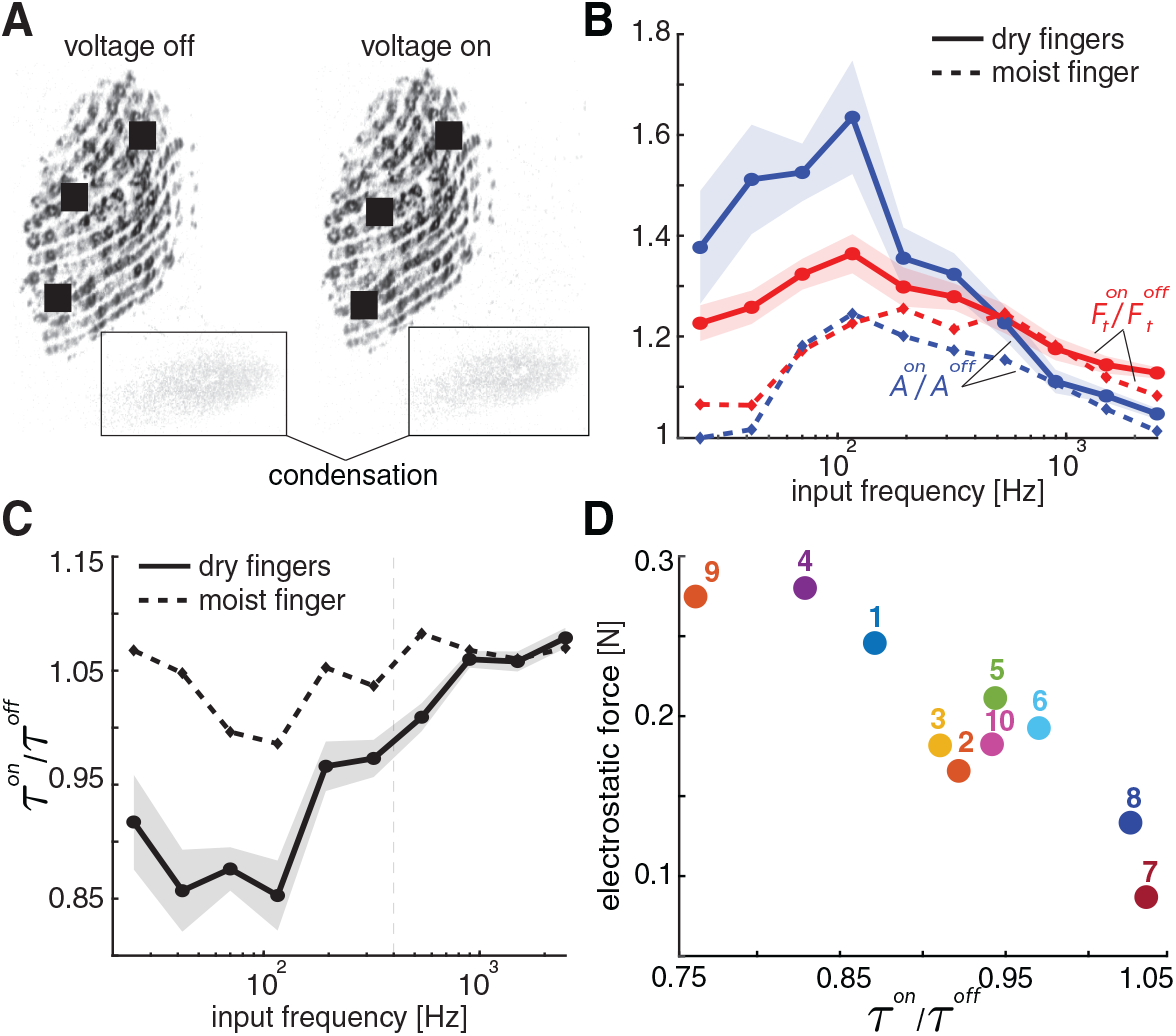
Effect of moisture on electrostatic actuation. (A) Representative fingerprint images showing condensation, indicating the presence of moisture. (B) Ratios of voltage on/off ratios values for real contact area and tangential force, and (C) interfacial shear stress comparing participants with moist and dry fingers. (D) Mean electrostatic force within the vibration regime for each participant. with drier fingers, whose images did not exhibit condensation (Fig. 4C). Within the vibration regime, participants with moist fingers exhibited a mean interfacial shear stress ratio that was 15.3% higher than those with dry fingers. When moist-finger data were excluded, the transition frequency shifted upward to 400 Hz. During this regime, participants with moist fingers showed lower electrostatic force and electrical impedance (Fig. 4D, Figs. S25–S27). Across participants, larger reductions in interfacial shear stress under voltage-on conditions were associated with higher electrostatic force, with a similar but weaker trend for electrical impedance (Fig. 4D).

## Discussion

Fingertip contact on electrostatic displays is shaped by oscillating electric fields that attract the fingertip toward the surface, thereby modifying the real contact area and friction at the skin–glass interface. By directly measuring time-resolved contact area and forces, we demonstrate that the frequency of electrostatic actuation governs fingertip–surface interactions, giving rise to two regimes: a *vibration* regime at low frequencies and an *adhesion* regime at high frequencies. These regimes produce characteristic changes in tangential force, real contact area, and interfacial shear stress. In the vibration regime, the increase in contact area exceeds the rise in tangential force between voltage-on and voltage-off conditions, resulting in a net reduction in interfacial shear stress. While electrostatic adhesion increases both friction and contact area regardless of frequency, at low frequencies, vibration weakens the adhesive contribution to tangential force, thereby reducing overall frictional enhancement. This vibration-induced friction reduction is consistent with prior observations across a range of materials and systems (*15, 17, 19, 27, 28*). In this regime, electrostatic actuation induces periodic normal force oscillations that diminish the effective mechanical coupling between the fingertip and the touchscreen, leading to a decrease in interfacial shear stress (*17*). According to the quasi-static model (*19*), these oscillations promote stick–slip dynamics at the interface, producing intermittent slipping and a reduction in effective tangential force—consistent with the displacement predicted by the spring-mass-damper model (Fig. S9). A similar reduction in the interfacial shear stress has been reported for finger–glass interaction under mechanical vibration (*17*), supporting the interpretation that vibration primarily reduces tangential force rather than contact area. Analysis of supplementary videos (Movies S1 & S2) further revealed clear within-cycle modulation of tangential motion at low frequencies, including pronounced (2 *f*_0_) components in instantaneous velocity (Fig. S28). This behavior is consistent with partial stick-slip motion, supported by corresponding within-cycle modulation in instantaneous interfacial shear stress (Fig. S29).

Notably, the observed peak around 116 Hz in contact area, tangential force, and interfacial shear stress ratio coincides with both the frequencies of enhanced tactile sensitivity of electrovibration (*13*) and the normal-direction resonant modes of the fingertip (*29*). This agreement suggests that the pronounced low-frequency vibratory effects arise from the combined influence of the electrical properties of the finger-display system and fingertip mechanics (Fig. S30).

At frequencies above 320 ± 82 Hz, the interfacial shear stress ratio between the voltage-on and voltage-off conditions exceeds unity, marking the transition to the adhesion regime. In this regime, the fingertip is less able to mechanically track rapid oscillations due to the constraints imposed by its intrinsic viscoelastic response (*14, 30*). Consistently, previous studies modeling finger–touchscreen interaction under electrostatic actuation as a spring-mass-damper system have also shown that the system behaves as a low-pass filter, with induced oscillations attenuating above 300 Hz (*14*). This behavior aligns with the interval-averaged differences observed here between the vibration- and adhesion-dominated regimes. As the vibration component of electrostatic actuation diminishes, adhesive effects become dominant, yielding interfacial shear stress above baseline (*τ*^on^/*τ*^off^ ≥ 1). Crucially, this adhesion-dominated regime does not reflect increased intrinsic adhesive strength; rather, it results from the skin’s limited ability to track high-frequency oscillations, reinforcing a continuous contact state. This physical transition aligns with qualitative feedback from participants, who described a distinct ‘stickier’ sensation in the adhesion regime compared to the ‘vibratory’ feel at lower frequencies.

Furthermore, the inverted U-shaped frequency response observed in electrostatic force, real contact area, and tangential force arises from the coupled mechanical and electrostatic dynamics at the interface. The vertical electrostatic attraction between the finger and surface increases the real contact area, which in turn enhances tangential force. A similar U-shaped trend in electrostatic force was reported in (*8, 22*), where the decrease at low frequencies was attributed to charge drift toward the outer surface of the stratum corneum (*6–8*). At high frequencies, the frequency-dependent dielectric properties of the stratum corneum limit sufficient charge migration within each oscillation cycle (*T* = 2*π*/*ω*), thereby reducing the effectiveness of electrostatic actuation. This interpretation is supported by electrical impedance measurements of the finger-touchscreen (*18*,*22*,*31*,*32*), which similarly showed a reduction at higher frequencies, consistent with reduced electrostatic force.

Across participants, the modulation of contact area, tangential force, and interfacial shear stress varied significantly, reflecting individual differences in skin moisture and electrical properties. Participants with moist fingers exhibited lower electrostatic force and electrical impedance, consistent with prior findings that skin hydration enhances conductivity and reduces the effective air gap, weakening the electric field at the interface (*32*). Although these individuals showed greater baseline friction and contact area without actuation (*24, 26, 32, 33*), they exhibited much smaller relative increases with electrostatic loading. This behavior can be attributed to greater effective damping in moist skin, which limits vertical displacement and reduces the modulation of tangential force and contact area. These findings indicate that increased skin moisture primarily suppresses the vibratory component of electrostatic actuation.

Beyond moisture, additional inter-participant differences in fingerpad curvature and skin mechanics likely contributed to observed variability. As shown in Fig. 3G and Figs. S22–S24, electrostatic actuation tended to increase contact area near the fingertip center, where vertical electrostatic force draws the skin toward the surface. The magnitude and spatial distribution of this expansion varied across participants. The transition frequency separating the vibration and adhesion regimes also differed across participants, further reflecting mechanical and geometric variability. Together, these electrical, mechanical, and physiological differences constrain the uniformity of electrostatic modulation of fingertip–surface interactions.

Despite careful investigation, this study has several limitations. First, the FTIR-based imaging and analysis method (*34*) provides an optically resolved measure of fingertip–glass real contact area and may not capture its absolute magnitude at the nanoscale. As real contact area is inherently scale-dependent, multiscale approaches combining optical and AFM measurements suggest that unresolved sub-optical roughness can influence its absolute value (*35*). Accordingly, this limitation may affect the absolute magnitude of the measured area; nonetheless, the present study focuses on relative changes measured consistently across conditions. Second, the dynamic bandwidth of the experimental setup may attenuate high-frequency oscillations. Consequently, the analysis primarily relies on interval-averaged metrics rather than fully resolved instantaneous oscillation amplitudes (see Supplementary Material and Fig. S31 for details). Finally, the inter-participant differences were inferred from image condensation, electrical impedance, and electrostatic force; direct measurements of skin hydration and other biomechanical properties were not performed. Incorporating such measurements in future work would provide a stronger foundation for interpreting inter-participant variability in electrostatic actuation.

From a haptic display design perspective, the present results identify distinct operating regimes for electrostatic interfaces under conditions comparable to those studied here. Frequencies in the vibration-dominated regime, particularly near the observed peak around 116 Hz, are well suited for enhancing dynamic contact modulation and vibratory tactile feedback. In contrast, frequencies above the transition near 320 Hz, where oscillatory skin deformation is attenuated and interfacial shear stress is restored or increased, are more appropriate for interactions intended to feel more adhesive or drag-like. The reduced modulation observed for moist fingers further indicates that practical devices may benefit from adaptive or user-dependent tuning rather than fixed actuation parameters.

In conclusion, our direct measurements of finger contact-area modulation under oscillating electric fields demonstrate that contact dynamics of sliding fingers on electrostatic displays are jointly governed by the interplay of vibration and adhesion, with vibratory effects diminishing at higher frequencies due to the electromechanical response of the finger-display system. These findings provide a mechanistic foundation for electrostatic haptic interfaces and support the development of more reliable, adaptive, and skin-condition-aware designs.

## Materials and Methods

The experiment was conducted with ten participants (seven men, three women; mean age: 27, SD:±2.45). The study was conducted by adhering the Declaration of Helsinki and approved by the Ethics Council of TU Delft (application no. 5108). All participants provided informed consent.

The participants slid their right-hand index finger across a capacitive touchscreen (SCT3250, 3M Inc.), which was electrostatically actuated by an alternating voltage applied to its conductive layer. The voltage signal was generated using a data acquisition card (PCIe 6321, NI Inc.) and amplified by a high-voltage amplifier (9200A, Tabor Electronics). Participants wore an anti-static wrist strap during the experiment. The touchscreen was mounted on two six-axis force sensors (Nano17 Titanium, ATI Inc.) to record contact forces at a sampling rate of 10 kHz. Finger motion was controlled by a motorized linear stage (NRT150/M, Thorlabs Inc.) set at a fixed angle of 60◦. Electrical impedance was measured via a differential probe and shunt resistor positioned between the amplifier and the touchscreen. Fingertip contact area was recorded using a high-speed camera at 1000 fps (MotionBLITZ EoSens mini2, Mikrotron) with a lens (LM16HC, Kowa) mounted beneath the glass, employing the FTIR method (*17*). The glass was illuminated using a light source (KL 2500 LED, Schott) fitted with a custom collimation package and diffuser. The touchscreen assembly was mounted on damped posts attached to an optical breadboard to minimize transmission of external vibrations. Data acquisition from the force sensors, linear stage control, and camera triggering were synchronized using MATLAB Simulink. Force and impedance data were collected through Simulink, while contact images were captured using MotionBLITZ software. Real contact area was computed from contact images following (*34*). See Supplementary Material for details of data extraction and analysis.

Before the experiment, participants were instructed to wash their hands and wipe them with a microfiber cloth. During each trial, the participant’s finger was moved laterally at a constant speed of 20 mm/s while maintaining a normal force of 1 N, guided by real-time LED feedback. Data were recorded only when the applied normal force was within ±10% of the target, and the fingerprint image was clearly visible. Electrostatic actuation was applied during the same sliding motion using alternating conditions: voltage-off and voltage-on at 100 V. Ten logarithmically spaced sine wave frequencies (25–2500 Hz) were tested, each with three repetitions. The full experimental session lasted approximately 30 minutes per participant. To minimize variability due to moisture, participants wiped their fingertips with a microfiber cloth before each trial, and a fan was used to maintain consistent skin dryness.

## Supporting information

Supplementary Information for: Electrostatic Actuation Induces Competing Adhesion and Vibration Regimes at Fingertip Contact

## Author contributions

CUK: Conceptualization, Methodology, Investigation, Software, Hardware, Formal Analysis, Data Curation, Visualization, Writing - original draft. MW: Conceptualization, Methodology, Formal Analysis, Visualization, Writing - review & editing, Supervision. YV: Conceptualization, Methodology, Formal Analysis, Visualization, Writing - original draft, Writing - review & editing, Supervision, Resources, Project Administration, Funding Acquisition.

## Data and materials availability

The data and code used in this paper will be publicly available upon acceptance

The authors used OpenAI’s ChatGPT and Grammarly Inc.’s Grammarly for language editing and readability improvements; the authors are responsible for all content and the final manuscript.

## Supplementary Information

Materials and Methods

Additional Results

Figures S1 to S31

Movies S1 to S5

