## Supplementary Information for: Electrostatic Actuation Induces Competing Adhesion and Vibration Regimes at Fingertip Contact for "Electrostatic Actuation Induces Competing Adhesion and Vibration Regimes at Fingertip Contact"

Celal Umut Kenanoglu[1], Michaël Wiertlewski[1], Yasemin Vardar[1*]
[1]Department of Cognitive Robotics, Delft University of Technology, Delft, 2628 CD, The Netherlands

#### Supplementary Text

##### Materials and Methods

###### Data Acquisition and Extraction

Fig. S1 illustrates the experimental setup; full details are provided in the Materials and Methods section of the main text. All control and data acquisition, except for image capture, were performed in MATLAB Simulink. Fingerprint images were recorded with a high-speed camera controlled by the MotionBLITZ software.

During the experiments, the fingertip of the participant was moved at a constant speed by a linear stage. The participant adjusted their normal force through the guidance of real-time LED feedback. Data were recorded only when the applied normal force remained within $\pm 10\%$ of the target and the fingerprint image was clearly visible. A camera trigger was sent when the finger entered the field of view. Only the tangential force data within this trigger window were used for analysis. In each trial, the voltage was initially off and then switched on by a second trigger, allowing voltage-off and voltage-on conditions to be compared within the same sliding motion.

Fig. S2a shows the raw tangential force data for all sliding directions (left-to-right and right-to-left). Fig. S2b shows the corresponding absolute tangential force during the camera trigger periods, combining all valid trials from Fig. S2a. An example close up from a single trial is presented in Fig. S2c, where the sinusoidal pattern of the tangential force is clearly visible. The minima of this waveform—corresponding to the zero-crossings of the voltage signal ($V = 0$ V)—closely match the tangential force values measured under the voltage-off condition, confirming consistency between the two cases.

When the camera trigger was sent from MATLAB Simulink to the MotionBLITZ software, the camera began recording at 1000 frames per second (fps). As with the tangential force measurements, we used the voltage-on trigger shown in Fig. S2a to extract data for both voltage-off and voltage-on conditions within the same trial. Raw fingerprint images could not be used directly due to optical distortion from lens curvature and the oblique viewing angle between the camera and the glass surface. First, radial distortion was corrected using intrinsic parameters obtained from checkerboard-based camera calibration. Next, a projective transformation [1] was applied to simulate a top-down view, as if the camera were orthogonal to the glass. To compute the transformation matrix, an image of a circular rubber piece placed on the glass was captured and the distorted ellipse was mapped onto a true circle. Real contact area was then extracted from the corrected images using the method in [2].

Fig. S3a shows the measured real contact area during the camera trigger period. An example from Trial 1 is shown in Fig. S3b, where the sinusoidal variation in contact area is clearly visible. The troughs of this waveform—corresponding to the zero-crossings of the voltage signal ($V = 0$ V)—match the real contact area values observed under voltage-off conditions, confirming consistency between the two states. As discussed in the main text, the increase in real contact area arises from the pulling effect of the electrostatic force.

The Fourier transform of the tangential force signal, presented in Fig. S5a, exhibits a prominent peak at twice the electrostatic actuation frequency, in agreement with previous observations by Vardar and Kuchenbecker [3]. This frequency doubling arises from the nonlinear dependence of the electrostatic force on the applied voltage. The electrostatic force can be expressed as [4]

$$F_e = \epsilon_0 \frac{AV^2}{2\left(\frac{d_{\text{ins}}}{\epsilon_{\text{ins}}} + \frac{d_{\text{air}}}{\epsilon_{\text{air}}} + \frac{d_{\text{SC}}}{\epsilon_{\text{SC}}}\right)^2 \epsilon_{\text{air}}}, \quad (1)$$

where $A$ is the contact area of the fingerpad; $d_{\text{ins}}$, $d_{\text{SC}}$, and $d_{\text{air}}$ denote the thicknesses of the touchscreen's insulating layer, the stratum corneum, and the air gap between the touchscreen and the finger, respectively; $\epsilon_{\text{ins}}$, $\epsilon_{\text{SC}}$, and $\epsilon_{\text{air}}$ are their relative permittivities; and $\epsilon_0$ is the permittivity of free space. As seen in Eq. 1, the electrostatic force depends on the contact area, finger properties, and the air gap during interaction, while the remaining parameters are fixed for a given touchscreen configuration. Because the electrostatic force scales with the square of the applied voltage, a sinusoidal voltage of the form $V = V_0 \sin(\omega t)$ with no DC offset yields

$$V^2 = V_0^2 \frac{1 - \cos(2\omega t)}{2}$$

following the trigonometric identity $\sin^2(\omega t) = \frac{1-\cos(2\omega t)}{2}$. This explicitly reveals a component at $2\omega$, and thus a frequency-doubling effect in the electrostatic force. The Fourier transform of the real contact area signal, presented in Fig. S5b, likewise exhibits a prominent peak at twice the electrostatic actuation frequency, consistent with the nonlinear voltage–force relationship described in Eq. 1 and similar to the behavior observed for the tangential force in Fig. S5a. Representative fingerprint image sequences showing the cyclic increase and decrease of real contact area in response to the applied electrostatic force are provided in the main text.

An example of the current signal from a single trial is shown in Fig. S4. In this figure, the DC offset has been removed, revealing a sinusoidal waveform whose peak occurs at the electrostatic actuation frequency. The Fourier transform of the current signal, presented in Fig. S5c, exhibits a prominent peak at the electrostatic actuation frequency, as expected for a signal that directly reflects the voltage drive applied to the touchscreen.

#### Implementation of the Spring-Damper-Mass Model

We model the fingertip normal dynamics as a linear spring-damper-mass system with displacement $u(t)$ under an electrostatic force input $F_e(t)$:

$$m\ddot{u}(t) + b\dot{u}(t) + k\,u(t) = F_e(t). \quad (2)$$

The complex frequency response is

$$H(\mathrm{j}\omega) = \frac{u(\mathrm{j}\omega)}{F_e(\mathrm{j}\omega)} = \frac{1}{k - m\omega^2 + \mathrm{j}b\omega}. \quad (3)$$

In our experiments, a sinusoidal voltage drive at the base frequency $f_0$ produces an effective electrostatic excitation at $2f_0$, so we evaluate the response at $\omega_{\text{eff}} = 2(2\pi f_0)$. The electrostatic force amplitude at each base frequency was obtained by averaging across participants; this served as the input $F_e$ for the single-degree-of-freedom model. $F_e$ was then estimated from the experimentally measured friction coefficients using $(1-\mu_{\text{off}}/\mu_{\text{on}})F_n$ [5, 6, 7]. The resulting displacement was then compared against contact-area modulation measured independently by the FTIR imaging. The steady-state displacement amplitude at the effective frequency was then computed from Eq. (3) as $|u(\omega_{\text{eff}})|$. The lumped parameters $k$, $b$, and $m$ were chosen within ranges reported for fingertip dynamics in prior studies [8, 9, 10, 11, 12]. As these parameters are known to vary across individuals and experimental conditions, they should not be interpreted as unique constants. For the lumped parameters, we used $k = 1500$ N/m, $b = 0.7$ N s/m, and $m = 1 \times 10^{-4}$ kg, which was selected as one representative parameter combination providing an approximate description of the experimentally observed frequency-dependent behavior. The sensitivity analyses of the parameter selection is presented in Fig. S11.

#### Implementation of Persson Contact Theory with Spring-Damper-Mass Model

In fingertip–glass contact, tissue compliance and surface roughness confine load to a subset of asperity peaks, so the true contact area is smaller than the apparent area. As attraction in the normal direction increases, additional micro-contacts form and existing ones grow, increasing the real area. The interfacial separation $u(x, y)$ varies laterally and is well approximated by a Gaussian distribution with standard deviation $u_{\text{rms}}$. Many engineering and biological surfaces are self-affine; their roughness depends on magnification. The glass counterface is treated as perfectly rigid and smooth.

After obtaining the displacement amplitude from the spring–mass–damper model (evaluated at the effective excitation $2f_0$), we map it to real area using a mean-field (Persson-type) area–separation relation [10]:

$$\frac{A^{\text{on}}}{A^{\text{off}}} = \exp\left(\frac{u_m}{u_{\text{rms}}}\right), \tag{4}$$

where we set $u_{\text{rms}} = 53\ \mu$m [13]. Here the spring–mass–damper block provides the *macroscopic* displacement amplitude $u = u_{\text{macro}}$ from the electrostatic force (all fingertip identification experiments were performed at the macroscopic scale). Because the fingertip interface is heterogeneous (partial asperity saturation, non-uniform pressure under the ellipsoidal fingertip geometry, residual air gaps), the microscale separation does not track the macroscopic approach one-to-one. To reflect this, we introduce a single phenomenological mapping (macro scale→micro scale) used in Eq. 4, where $u_{\text{m}} = \gamma\, u$, meaning that only a fraction $\gamma$ of the macroscopic approach effectively changes the local gap at the microscale. Substituting $u_{\text{m}}$ in Eq. 4 prevents over-prediction of $A^{\text{on}}/A^{\text{off}}$ while keeping the SMD parameters within literature ranges. In our implementation, $\gamma$ is calibrated by least-squares fit to the measured area ratio and equals 0.161.

#### Implementation of the Quasi-static Friction Model

We also modeled the reduction in tangential force under vibration using a quasi-static framework [14], which attributes the decrease to stick-slip behavior. This model builds on Amontons' law of friction and incorporates contact stiffness in both the normal and tangential directions, enabling it to account for frictional effects under oscillatory loading conditions during sliding with constant velocity. It provides a macroscopic and physically interpretable framework for examining how normal oscillation can influence tangential frictional response. In this study, we adapted the quasi-static framework to finger–surface electrostatic interactions, where it has not previously been applied, to examine whether it can reproduce the experimentally observed frequency-dependent tangential forces.

In this work, we used this model as a first-order macroscopic approximation. Its use is motivated by the fact that electrostatic actuation introduces normal-direction oscillation, which is the type of input for which the model framework was developed. However, the model does not explicitly capture viscoelastic dissipation or other electrical contributions at the interface. It is therefore most useful here as a physically interpretable framework for examining whether vibration-induced normal oscillation can explain the observed difference between real contact area modulation and tangential force modulation, rather than as a complete dynamic predictive model.

The effective friction coefficient $\overline{\mu}$ under vibration by the quasi-static model can be written as [14]

$$\overline{\mu} = \mu_0(1 - \alpha\psi_w(\beta)) \tag{5}$$

where $\mu_0$ is the friction coefficient in the absence of oscillation (full slip), $\alpha$ is the dimensionless amplitude of oscillation, and $\beta$ is the dimensionless velocity.

$$\alpha = \frac{A_z}{d_z} \tag{6}$$

where $A_z$ is the amplitude of the normal displacement oscillation and $d_z$ is the mean indentation.

$$\beta = \frac{k_y v_0}{\mu_0 k_z A_z f} \tag{7}$$

where $k_y$ is the tangential stiffness, $v_0$, is the sliding velocity, $k_z$ is the normal stiffness, and $f$ is the effective frequency.

$$\psi_w = (1 - \beta/\beta_c)^{2.4} \tag{8}$$

is the equation of the $\psi_w$ under the harmonic oscillation. $\beta_c$ is the maximal positive value of its first derivative.

In our scenario, we observe a mismatch between the ratio of contact area and tangential force under normal oscillation. If normal oscillation (electrovibration) were absent, these ratios would likely align, reflecting the isolated effect of electroadhesion. Therefore, the observed divergence suggests that normal oscillation introduces an additional modulation mechanism beyond electroadhesion, likely contributing to enhanced frictional variation through dynamic contact changes. Therefore, if there is no normal oscillation, $\mu_0$ can be found as

$$\mu_0 = \mu_{off}\frac{A^{on}}{A^{off}} \tag{9}$$

Assuming adhesion scales friction with real contact area when normal oscillation is absent. The amplitude of the normal displacement occurs due to the electrostatic force, so we can find $\alpha$ as

$$\alpha = \frac{u}{F_n/k_z} \tag{10}$$

where $F_n$ is the normal force and $u$ ($A_z$) is the displacement due to electrostatic force from spring-damper-mass fingertip model. Here, we do not need any mapping factor because quasi-static model is already in macroscale model [14]. We can find $\beta$ as

$$\beta = \frac{k_y v_0}{\mu_0 k_z u(2f_0)} \tag{11}$$

When we substitute Eq. 8–11 to Eq. 5, we can find the reduced friction coefficient due to vibration under electrostatic force. We used our experimental data for all parameters except the stiffness of the finger in the vertical and lateral directions. Tangential stiffness is set in the range of [8, 9, 10, 11, 12] as $k_y = 900$ N/m, and same normal stiffness with spring-damper-mass model.

#### Additional Results

##### Frequency-dependent response of tangential force, real area, and interfacial shear stress

Figs. S6, S7, and S14 present box plots of the ratios of tangential force, real contact area, and interfacial shear stress between voltage-on and voltage-off conditions. All three measures exhibit a U-shaped dependence on frequency, consistent with the electrostatic force behavior shown in Fig. 3F of the main text and with previous studies [6, 7]. The variability in these ratios is notably higher in the vibration regime, where electrovibration is active, and this spread primarily reflects inter-participant differences. Fig. S8 shows the ratios of tangential force and real contact area together, explicitly highlighting the frequency range where the tangential force ratio exceeds the real area ratio.

##### Modeling of the modulation of real area and tangential force

Fig. S9 presents the displacement data obtained from the spring–damper–mass model. Similar to the $A^{on}/A^{off}$ ratio, it exhibits a pronounced peak at 116 Hz, followed by a gradual decline. In the adhesion regime, the displacement converges to zero, as the fingertip is less able to follow the rapid oscillations [9, 12].

We compared the contact area prediction of our combined spring-damper-mass and Persson contact theory with the prediction made by the mean-field contact theory under electroadhesion proposed in the literature. We followed the same procedure to estimate the contact area using mean-field theory, as described in [15, 16]. Fig. S10 compares the measured $A^{on}/A^{off}$ ratio with the predicted ratios using combined spring–damper–mass model with Persson's contact theory and Persson's contact theory under electroadhesion. Both models show good agreement with the experimental data. The combined spring–damper–mass and Persson model accurately captures the experimental results across all frequencies, with relative errors between 0.16% and 9.45% (mean absolute error = 4.7%). Similarly, Persson's contact theory under electroadhesion predicts the experimental data well, with relative errors ranging from 0.5% to 9% (mean absolute error = 4.2%). The spring–damper–mass with Persson model successfully reproduces the contact-area behavior in the vibration regime, where accounting for the fingertip's mechanical response is essential because the finger can follow the oscillations. In contrast, Persson's contact theory under electroadhesion more accurately represents the adhesion regime, as it excludes the effects of the fingertip's mechanical resonance and considers only the contribution of adhesion. In our analysis of Persson's contact theory under electroadhesion [15, 16], the thicknesses of the touchscreen insulating layer and the stratum corneum were set to $d_{ins}$=1$\mu$m and $d_{\mathrm{SC}}$=300$\mu$m, respectively, with a touchscreen dielectric constant of $\epsilon_{ins}$=8. The frequency-dependent dielectric function of the stratum corneum $\epsilon_{\mathrm{SC}}(\omega)$ was calculated using resistivity and permittivity values reported in [17]. In our analysis, we also consider that at low frequencies, charges on the skin surface may drift onto the touchscreen. This effect depends on the surface and bulk conductivity of the touchscreen, as well as the presence of conductive liquids (e.g., sweat or skin oils) in parts of the non-contact area. To account for this, we use a resistivity of $10^7\Omega$·m at low frequencies, consistent with [15, 16]. A semi-wet stratum corneum was assumed with a Young's modulus of $E$ = 20 MPa. The apparent contact area was set to $A_0$ = 43 mm$^2$, based on the average across participants as extracted from our fingerprint images. The overall procedure follows that described in [15, 16].

To assess model robustness, we performed a local sensitivity analysis by varying each lumped parameter by 20% around its nominal value while keeping the other two fixed (Fig. S11). The model retained the same overall frequency-dependent behavior under these perturbations. The mean fitting error changed from 4.59% to 5.12% for variations in $m$, from 5.17% to 5.07% for variations in $b$, and from 7.96% to 4.72% for variations in $k$, relative to 4.77% for the nominal case. These results indicate that the interpretation does not rely on a unique parameter set, although the response is more sensitive to stiffness than to mass or damping.

Fig. S12 presents the range of tangential stiffness values evaluated in our study. The quasi-static model shows good agreement with our experimental data in the vibration regime. When $k_y = 450$ N/m, the model errors range from 2.2% to 8.6%, with a mean error of 4.8%. For $k_y = 900$N/m, errors range from 1.5% to 8.5%, with a mean of 4.9%. When $k_y = 1800$N/m, the error ranges from 2.1% to 14.4%, with a mean of 5.4%.

Fig. S13 shows the friction reduction due to electrovibration, calculated using the quasi-static model when $k_y$ is selected as 900 N/m, as in the main text. This reduction is expressed as the ratio $1 - \bar{\mu}/\mu_0$, representing the friction coefficient when both vibration and adhesion are present, relative to when only electroadhesion is present (absence of oscillation). Similar to all measured signals, the curve in Fig. S13 exhibits a peak around 116 Hz.

**Representative measurements and statistical comparisons across regimes**

Representative tangential and real area measurements for all frequencies when the voltage is off and on are presented in Figs. S15 and S16. Fig. S17 shows the real contact area and tangential force data from both the vibration and adhesion regimes. In the vibration regime, both quantities exhibit oscillatory behavior, whereas in the adhesion regime the oscillations are attenuated, as the finger acts as a low-pass filter and is less able to follow rapid oscillations [9, 12]. Representative videos of the fingertip contact during the vibration and adhesion regimes are provided in Movies S1–S4. Note that no correction (i.e., optical distortion and angle correction) was applied to these movies to avoid making them identifiable. To further examine force-signal modulation, we compared low-frequency responses from two participants at 70 Hz (Fig. S18). In one participant, the tangential-force signal showed a clearly visible oscillatory component during electrostatic actuation, whereas in the other participant, the oscillatory component was markedly weaker. As this difference is observed at the same low actuation frequency, it suggests that the appearance of the raw force waveform is also influenced by subject-dependent fingertip dynamics and contact conditions.

Fig. S19 presents the results of post hoc paired t-tests on interfacial shear stress ratios across frequencies. To account for multiple comparisons, p-values were adjusted using the Benjamini–Hochberg procedure [18], which controls the false discovery rate while maintaining statistical power. These tests revealed significant differences between frequencies in the vibration and adhesion regimes ($p < 0.05$), highlighting a distinct behavioral transition. Notably, the contrast between the 70–116 Hz range and the adhesion regime was highly significant ($p < 0.001$). Fig. S20 and Fig. S21 show the corresponding post hoc paired t-tests with Benjamini–Hochberg correction for real contact area and tangential force across frequencies, respectively. These analyses also revealed significant differences between frequencies in the vibration and adhesion regimes ($p < 0.05$), with p-values for comparisons between the 70–116 Hz range and the adhesion regime being particularly low.

**Spatial distribution of contact and gap variation**

Fig. S22 presents raw contact images and normalized intensity from one participant. The intensity of the black regions—indicating real contact—is consistently greater at 100 V than at 0 V across all frequencies. The real contact area at 100 V is larger in the vibration regime than in the adhesion regime. To further assess the spatial distribution of contact, we visualized the fingerprint patterns using colormaps. These visualizations indicate that electrostatic actuation primarily increases contact in the central region of the fingertip, suggesting that the skin deforms more in this area under electrostatic actuation.

To further examine the spatial distribution of the contact increase under electrostatic actuation, we performed a spatial analysis by separating the fingertip contact region into central and peripheral zones of equal area. The central region was defined as the inner ellipse containing 50% of the fitted contact-ellipse area, and the peripheral region as the remaining outer 50%. For each region, we computed the ratio of real contact area at $V = 100$ to that at $V = 0$ from representative contact maps for one participant. For each frequency, the $V = 0$ and $V = 100$ contact maps were normalized together using a common maximum intensity value obtained from the combined pair of maps. Thus, the two maps at a given frequency share the same color scale and can be compared directly within that frequency. The normalization was performed separately for each frequency and was not applied globally across all frequencies. This analysis showed that the relative increase in contact area was generally larger in the central region than in the peripheral region. The ratio of real contact area at $V = 100$ to that at $V = 0$ in the vibration regime was 3.26 in the central region and 2.93 in the peripheral region. In the adhesion regime, the corresponding values were 1.46 and 1.38. These results indicate that the central enhancement was more evident in the vibration regime and became smaller in the adhesion regime. Representative spatial maps are presented in Fig. S23.

In addition, to highlight how the interfacial gap varies from the center toward the edges of the finger, we extracted a one-dimensional spatial gap profile from the fingerprint images for one participant (Fig. S24). A horizontal rectangular region with a height of 1 mm was placed across the middle of each fingerprint image, and the pixel values within this region were averaged column-wise from left to right. This procedure was performed separately for the voltage-on and voltage-off conditions, corresponding to $V = 100$ and $V = 0$, respectively. As the image intensity does not provide a directly calibrated physical measure of the gap, the resulting profile was treated as a relative gap proxy. For each frequency, the voltage-on and voltage-off profiles were normalized within that frequency before comparison. In this representation, lower values indicate a smaller relative gap, whereas higher values indicate a larger relative gap. The normalized profiles were then averaged within the vibration and adhesion regimes to obtain the mean spatial gap distributions for the two regimes. Overall, the difference between $V = 100$ and $V = 0$ remains relatively small near the edges of the finger contact, whereas it becomes more pronounced toward the center.

**Effects of fingertip moisture**

We observed the effect of moisture in some fingerprint images as visible condensation behind the finger, where the reduction in interfacial shear stress under electrostatic actuation was less pronounced. This condensation was evident in two participants, as shown in Fig. 4A (main text) and Movie S5. Electrostatic force and electrical impedance in the vibration regime were both lower for these participants compared to the rest (Fig. 4D (main text) and Fig. S25). Mean electrical impedance and electrostatic force in the adhesion regime are shown in Figs. S26 and S27. In this regime, the differences between moist and dry fingers in both electrostatic force and electrical impedance become less pronounced, and the interfacial shear stress under voltage-on conditions is slightly higher than under voltage-off conditions.

**Cycle-resolved response and interpretation of the 116 Hz peak**

To further examine the low-frequency frictional response, we performed an additional tracking analysis of fingertip motion from the supplementary video cases (Movies S1 and S2). Instantaneous tangential displacement and velocity were extracted over time for the tracked fingertip region. The resulting traces showed clear within-cycle modulation of tangential motion, consistent with partial stick-slip behavior during electrostatic actuation (Fig. S28). In both cases, frequency analysis revealed a pronounced peak at the effective actuation frequency ($2f_0$) in the tangential velocity signal. This indicates that the tangential motion was dynamically modulated within each electrovibration cycle. These cycle-resolved results provide further insight into the low-frequency regime and complement the cycle-averaged analysis of contact area, tangential force, and interfacial shear stress presented in the main manuscript. In addition, a time-resolved interfacial shear-stress estimate obtained from the measured tangential force and real contact area for one representative participant also showed clear within-cycle modulation, consistent with the tracked tangential-motion analysis (Fig. S29).

To better interpret the observed 116 Hz peak, we included a qualitative comparison based on the same spring–mass–damper model under three forcing assumptions. In the mechanical-only case, the electrostatic force amplitude was assumed to be constant across frequency. In the electrically weighted case, the same mechanical model was retained, but the force amplitude was modulated by an illustrative first-order high-pass-type weighting to represent a simple frequency-dependent electrical contribution [6, 12, 19]. In the third case, the model was driven by the experimentally inferred $F_e$. This comparison illustrates that the apparent peak location depends on how the forcing term is represented, and therefore supports interpreting the observed 116 Hz peak as part of a coupled electrical–mechanical fingertip–surface response rather than as a purely mechanical resonance alone (Fig. S30).

**Comparison with an auxiliary verification setup**

Fig. S31 provides additional verification of the dynamic behavior of the experimental setup. The figure compares an auxiliary verification setup, which used the same dual ATI Nano17 Titanium sensor arrangement as the main experiment but without the imaging subsystem, with the full experimental setup. Representative measurements from both setups were obtained from the same participant under fixed conditions. Figs. S31(a)–(c) present the impact-test result and representative tangential-force traces from the experimental setup at $f_0 = 100$ Hz and $f_0 = 600$ Hz, whereas Figs. S31(d)–(f) show the corresponding results from the verification setup at $f_0 = 100$ Hz and $f_0 = 600$ Hz. Although the two setups differ in their high-frequency impact-test response, both exhibit a clear tangential-force shift between the voltage-on and voltage-off intervals at higher frequencies. At lower frequencies, the minima of the tangential-force oscillation during the voltage-on interval remain close to the voltage-off level in both setups.

### Supplementary Figures

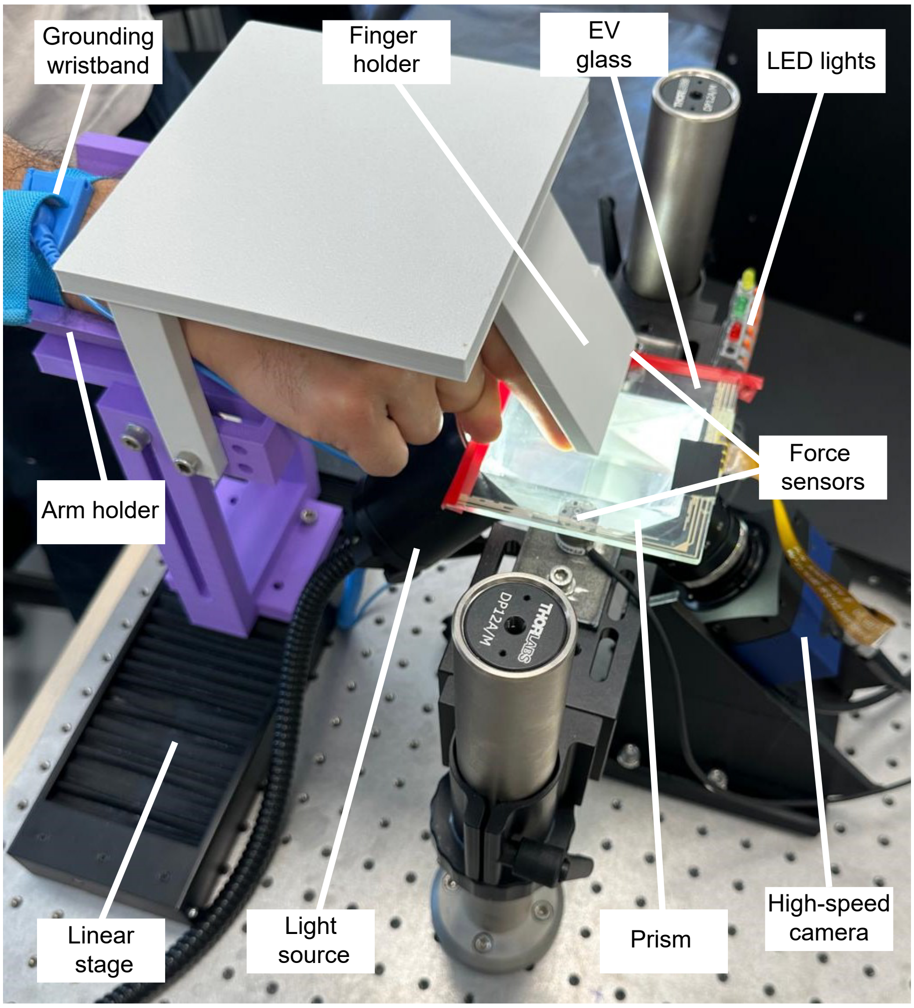


**Figure S1: Experimental setup.**

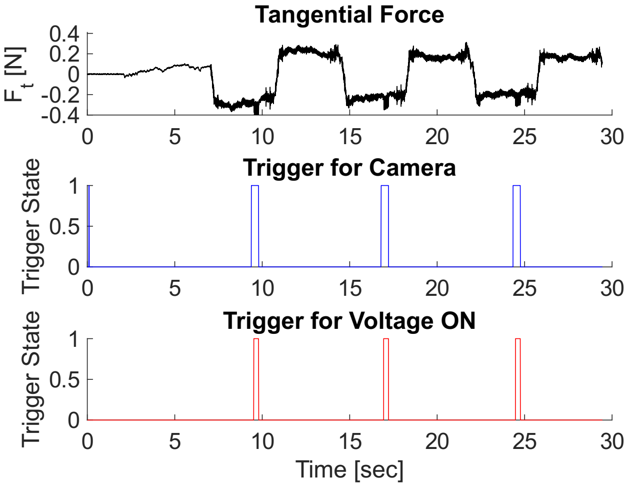


(a)

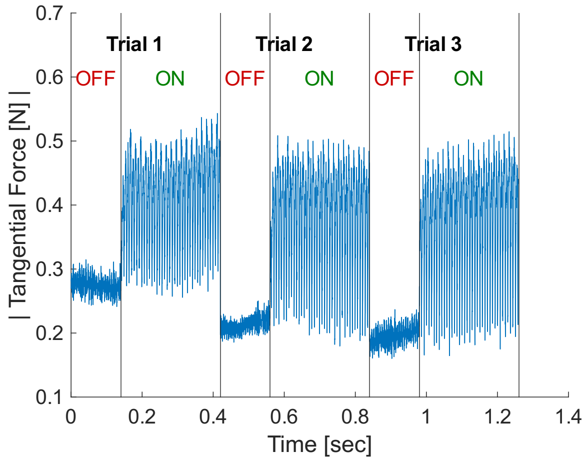


(b)

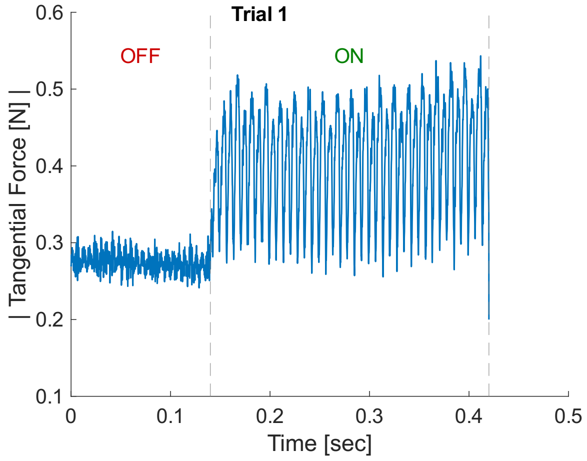


(c)

**Figure S2: Tangential force measurement.** Extracted tangential force and trigger signals; electrostatic actuation frequency is 70 Hz. (a) Raw tangential force data recorded during a trial with voltage and camera trigger signals. Value of 1 indicates trigger is active and value of 0 indicates trigger is not active. (b) Absolute value of tangential force data when camera trigger is on. (c) Absolute value of tangential force measurement for Trial 1.

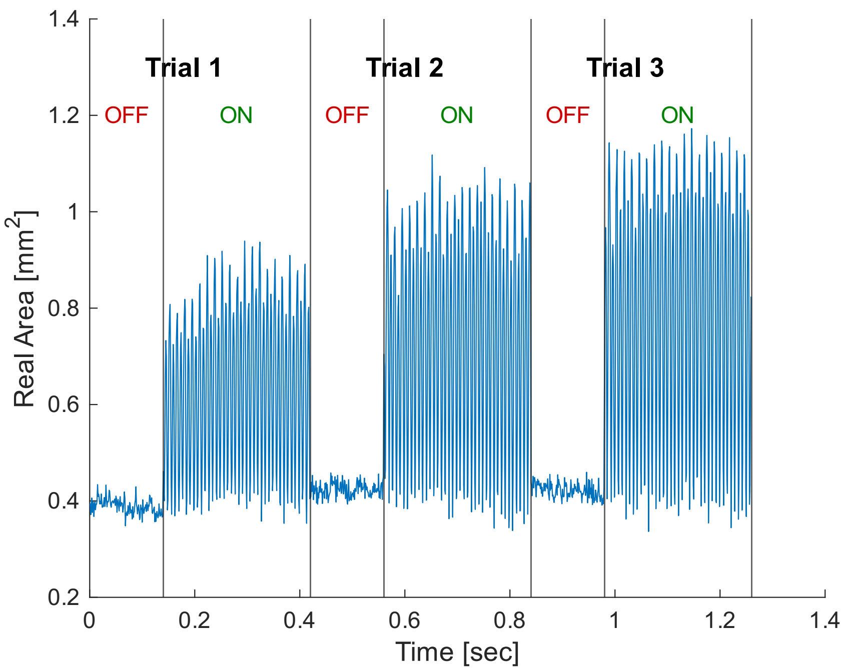


(a)

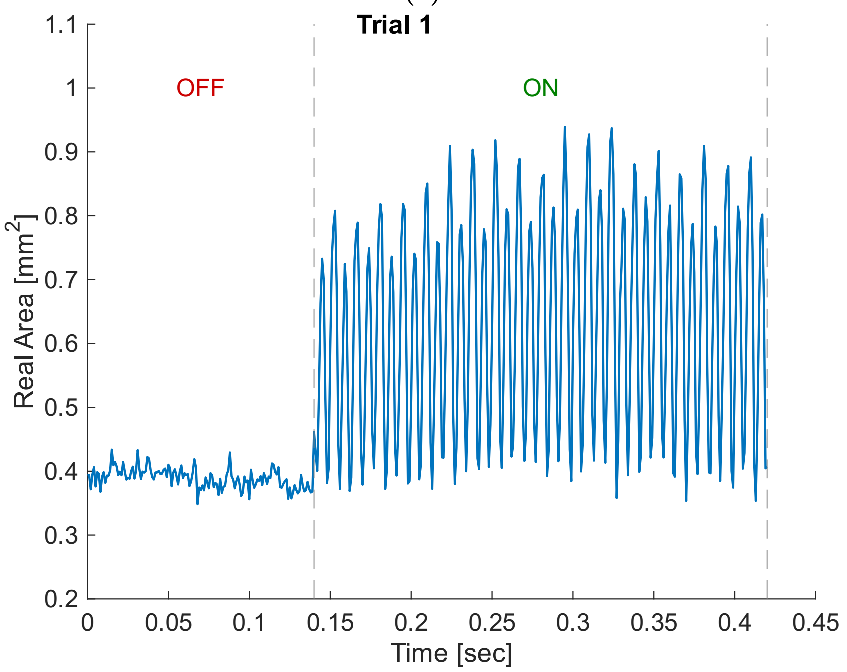


(b)

**Figure S3: Real area measurement.** Extracted real area data from fingerprint images; electrostatic actuation frequency is 70 Hz. (a) Real area data when camera trigger is on (All 3 trials for Voltage off and on). (b) Real area measurement for trial-1 when voltage off and on

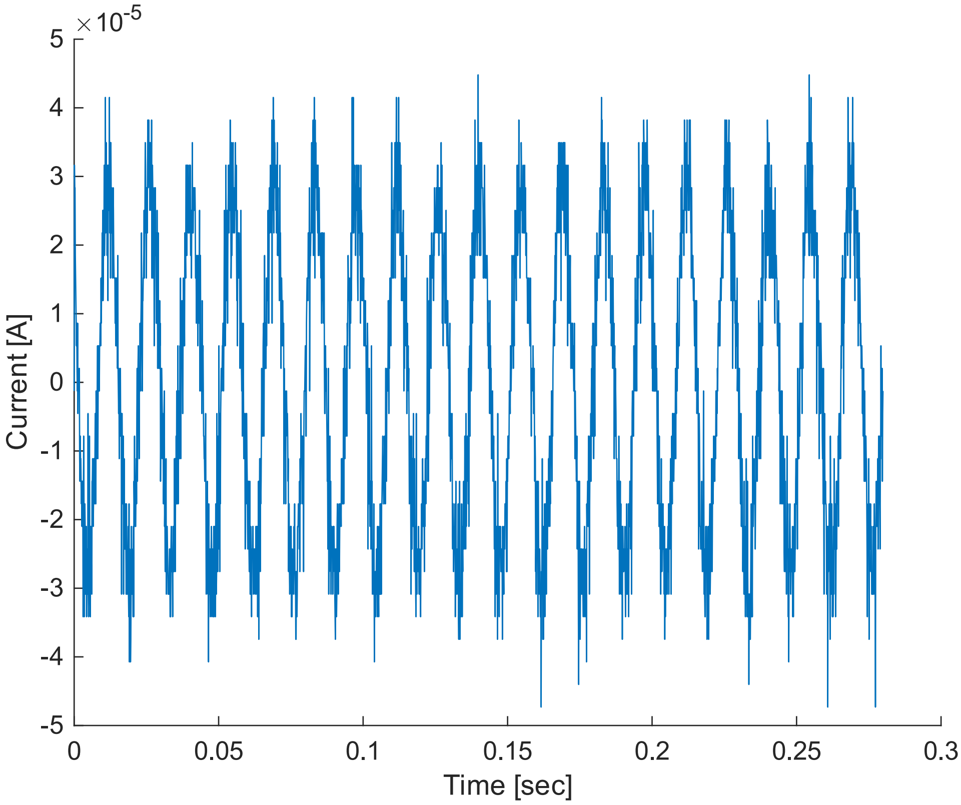


**Figure S4: Current measurement.** Example of current data recorded during Trial 1 when camera and voltage-on trigger are on; electrostatic actuation frequency is 70 Hz.

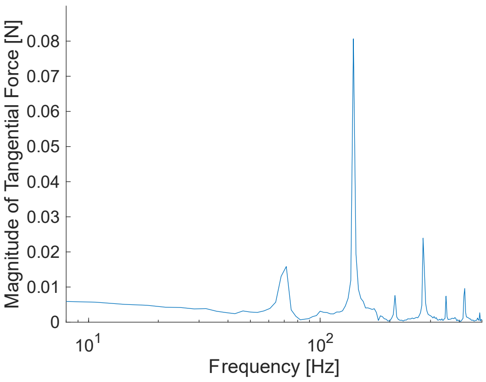


**(a)**

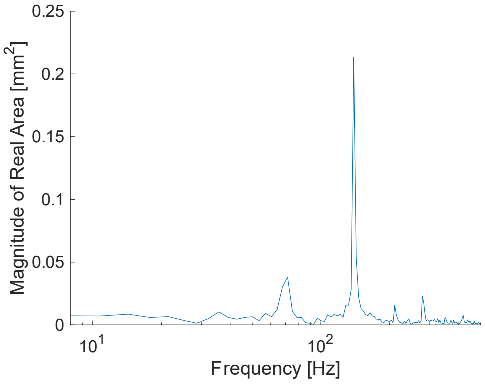


**(b)**

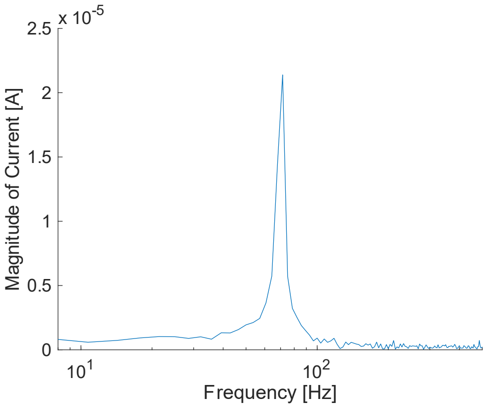


**(c)**

**Figure S5: Fourier transform of the recorded signals.** Fourier transform of (a) tangential force, (b) real area, and (c) current data during a trial with electrostatic actuation ($f_0 = 70$ Hz).

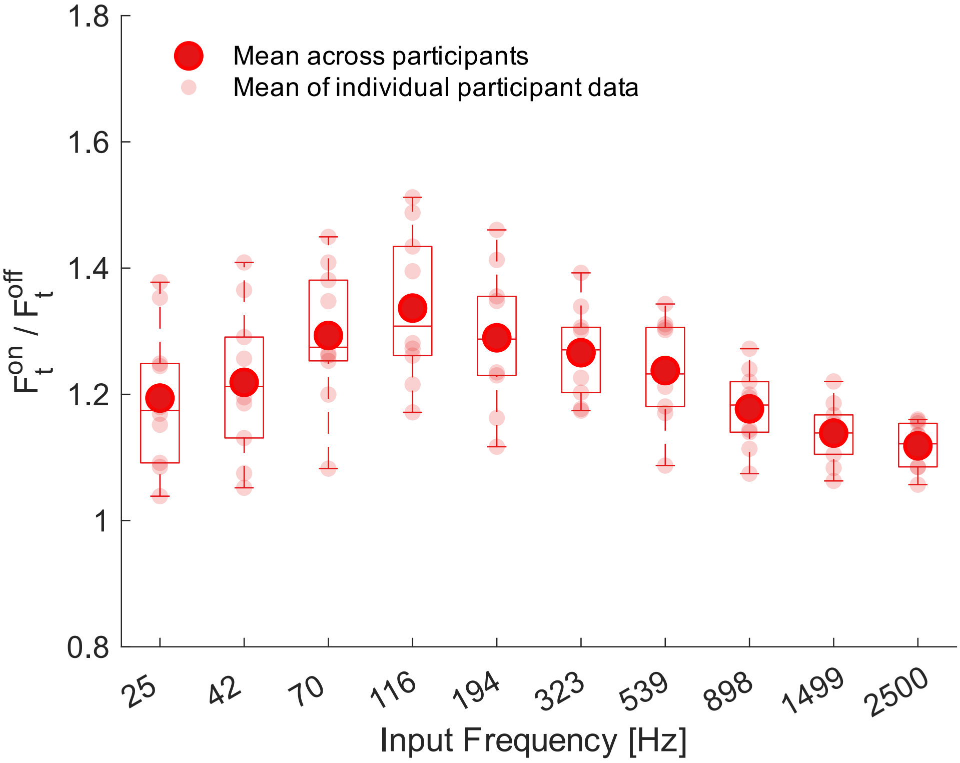


**Figure S6: Box plots of tangential force ratio.** Box plots of tangential force ratio condition when voltage on and off across frequencies. Large circles indicate group means; smaller faded circles represent the average of individual participants' repetitions.

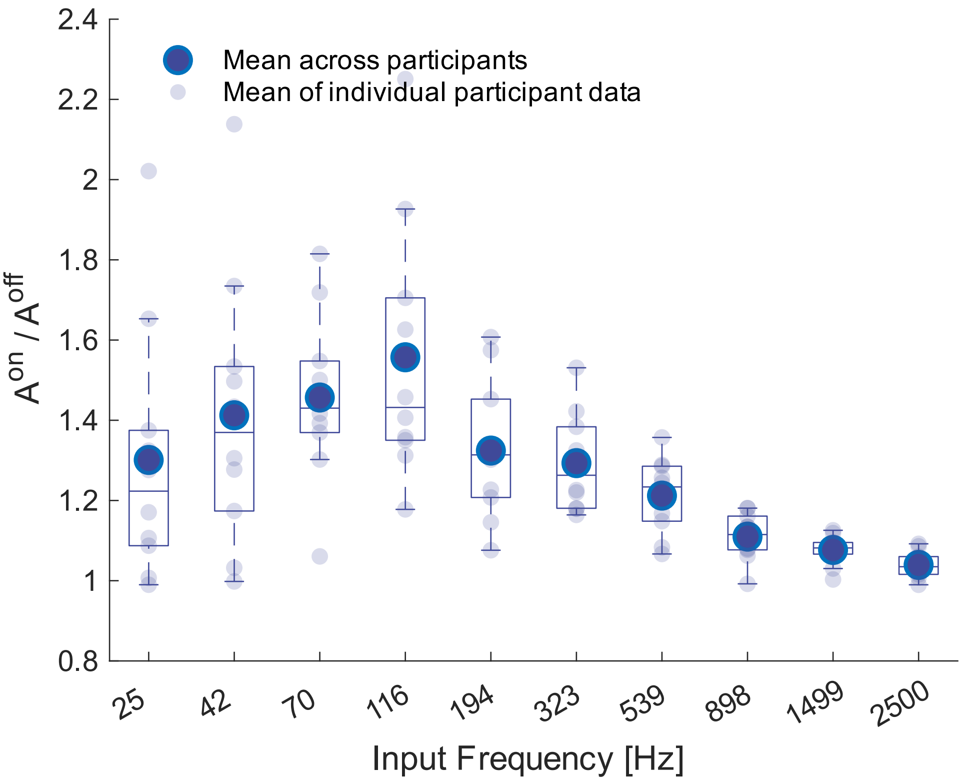


**Figure S7: Box plots of real area ratio.** Box plots of real area ratio condition when voltage on and off across frequencies. Large circles indicate group means; smaller faded circles represent the average of individual participants' repetitions.

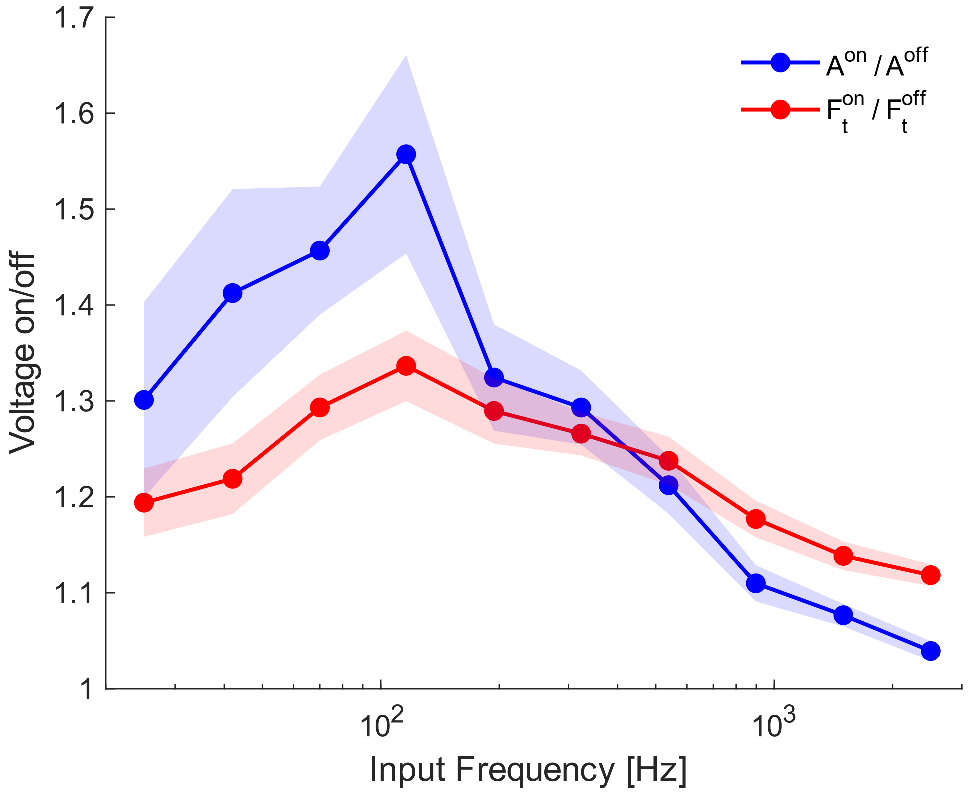


**Figure S8: Measured tangential force and real area ratio.** Real area and tangential force ratio when the voltage is on and off.

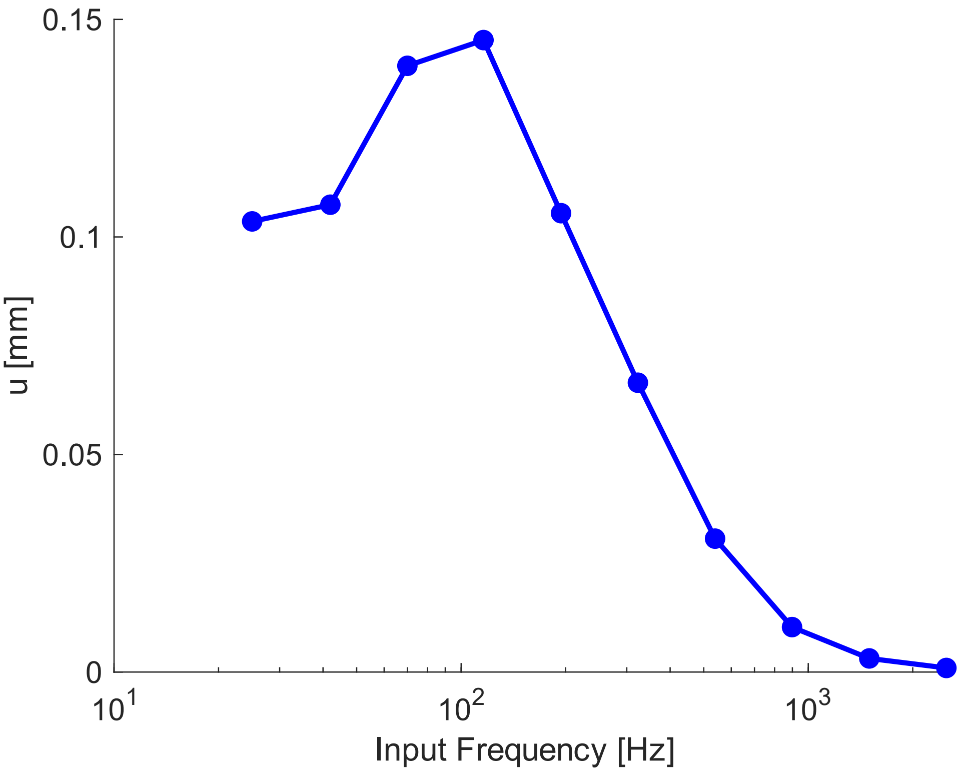


**Figure S9: Fingertip vertical displacement.** Finger displacement in the normal direction predicted with the spring-damper-mass model.

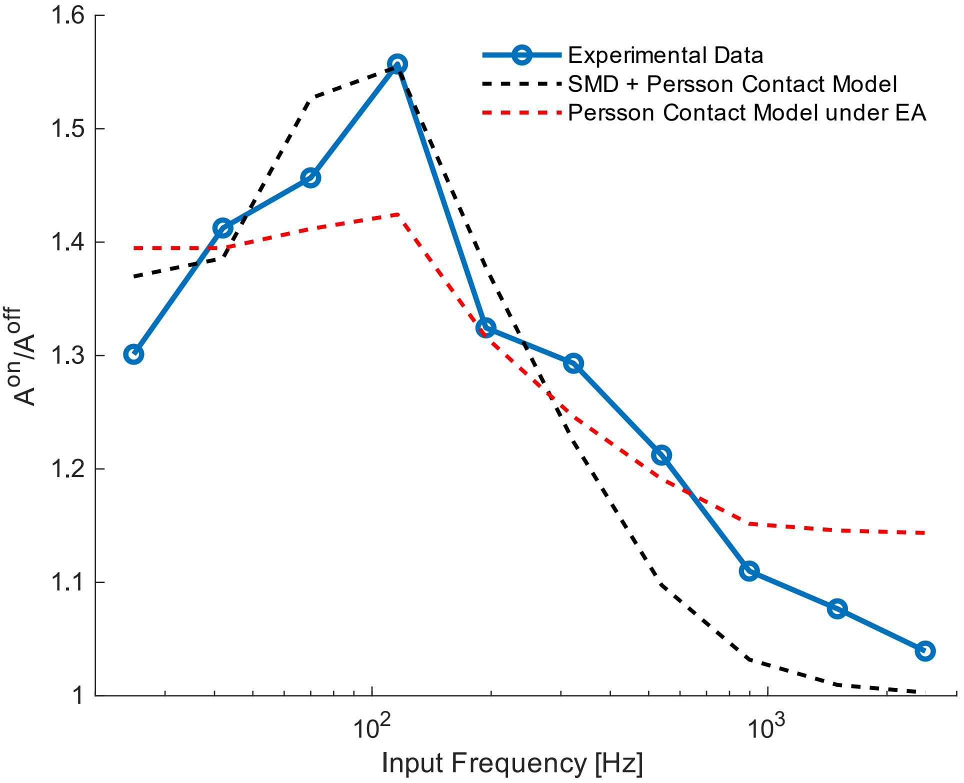


**Figure S10: Measured real contact area ratio with contact models.** Measured ratio of contact area ($A^{\mathrm{on}}/A^{\mathrm{off}}$) compared with predictions from the combined spring-mass-damper (SMD) and Persson contact models [10] and the Persson contact model under electroadhesion (EA) [15, 16].

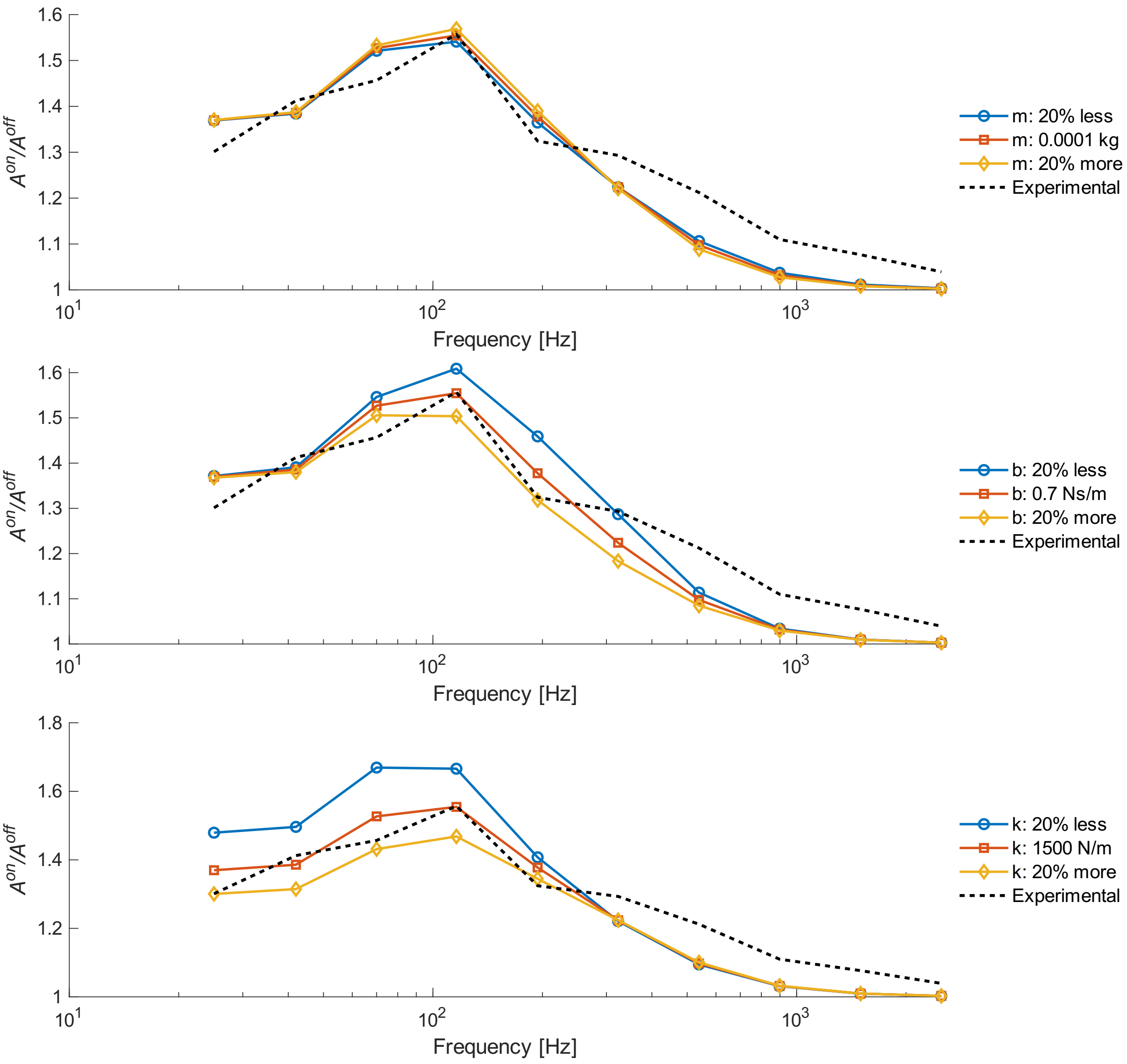


**Figure S11: Sensitivity analysis of selected lumped parameters.** Effect of ±20% changes in the model parameters mass (m), damping (b), and stiffness (k) on the predicted contact area ratio $A^{on}/A^{off}$.

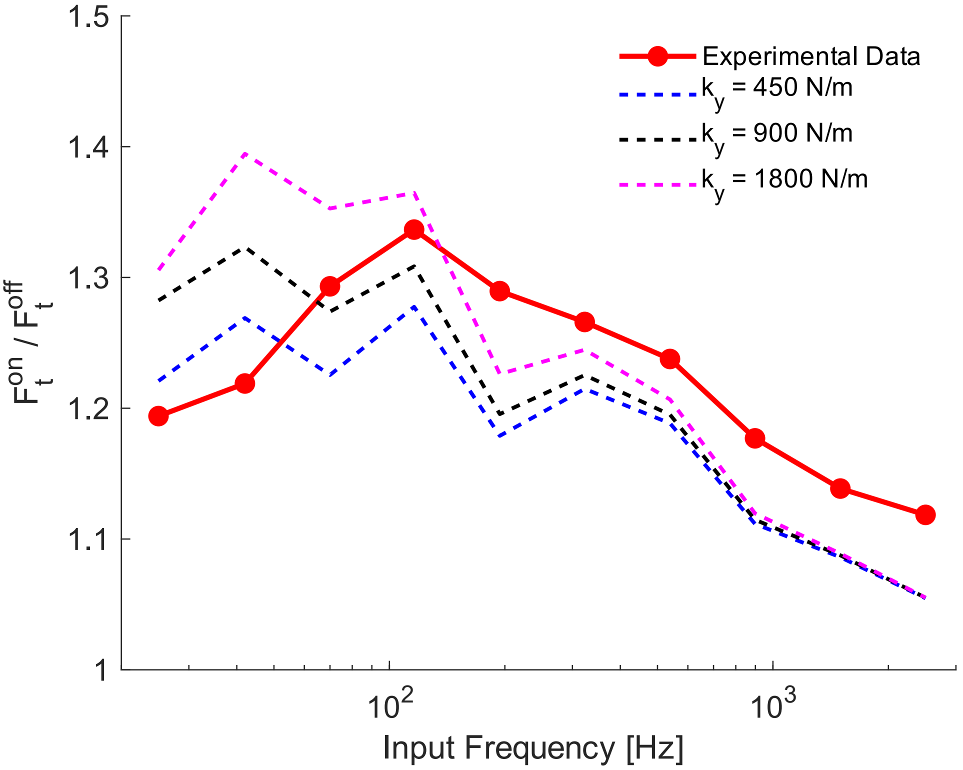


**Figure S12: Quasi-static model with various finger stiffness.** Results of the quasi-static model for three different finger stiffnesses.

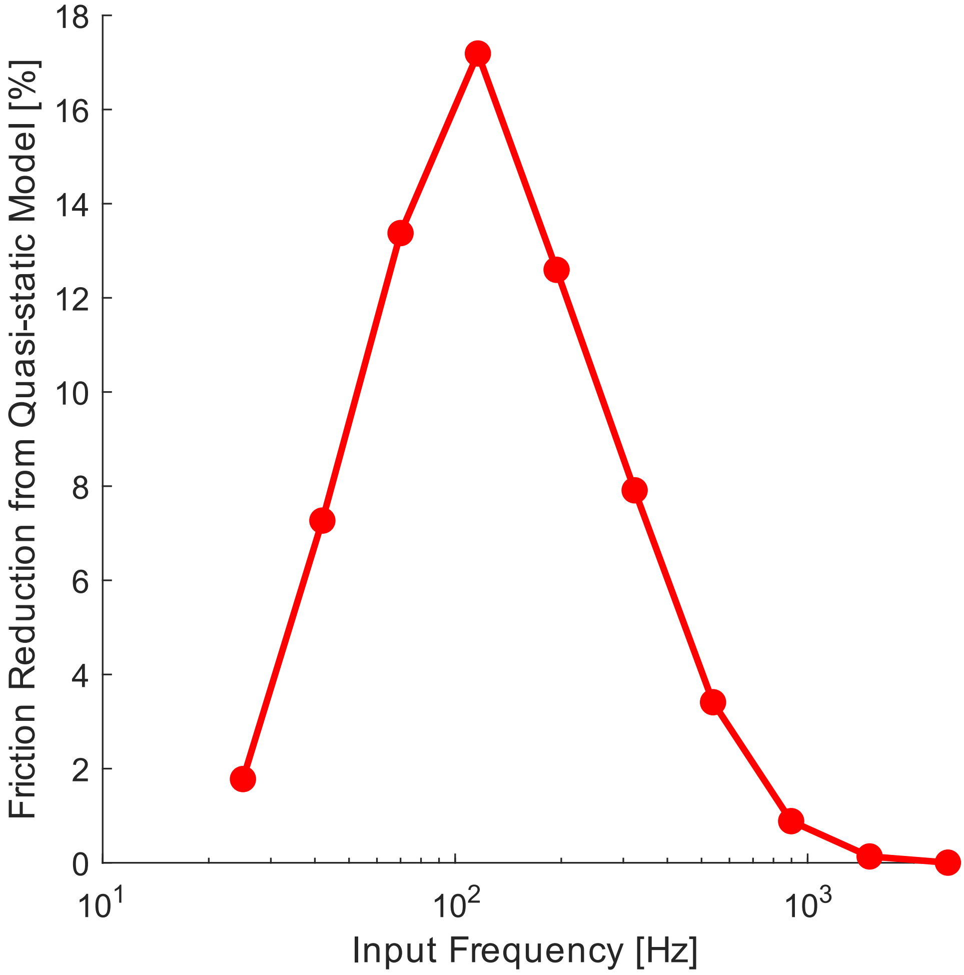


**Figure S13: Friction reduction from quasi-static model.** Friction reduction across frequencies calculated from quasi-static model.

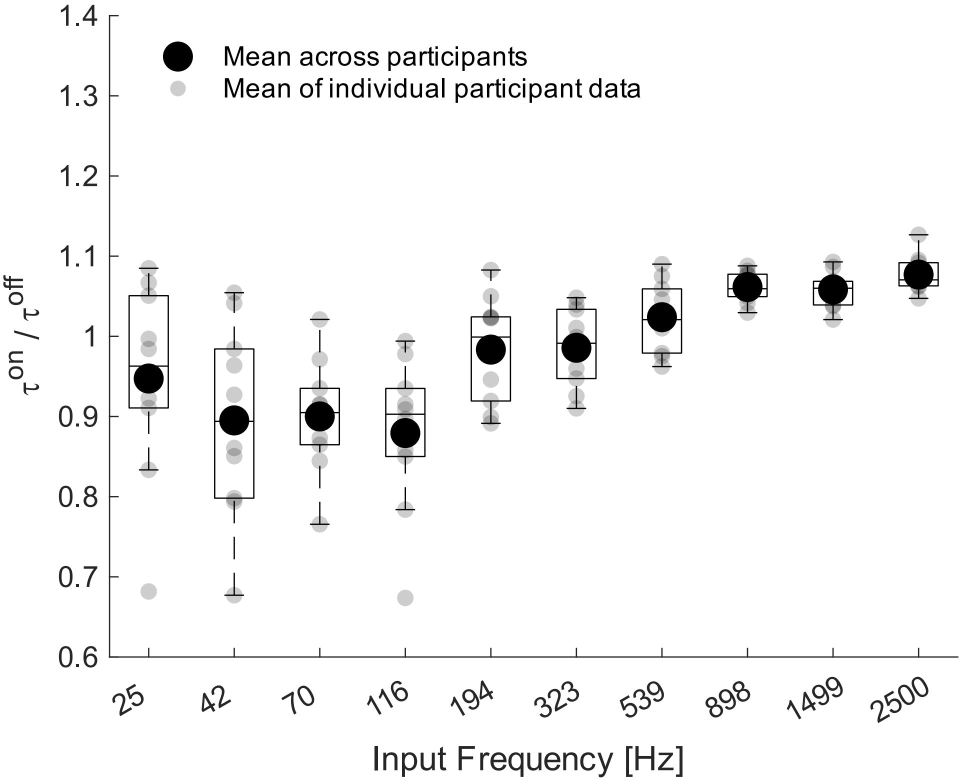


**Figure S14: Box plots of real area ratio.** Box plots of interfacial shear stress ratio ($\tau^{on}/\tau^{off}$) condition when voltage on and off across frequencies. Large circles indicate group means; smaller faded circles represent the average of individual participants' repetitions.

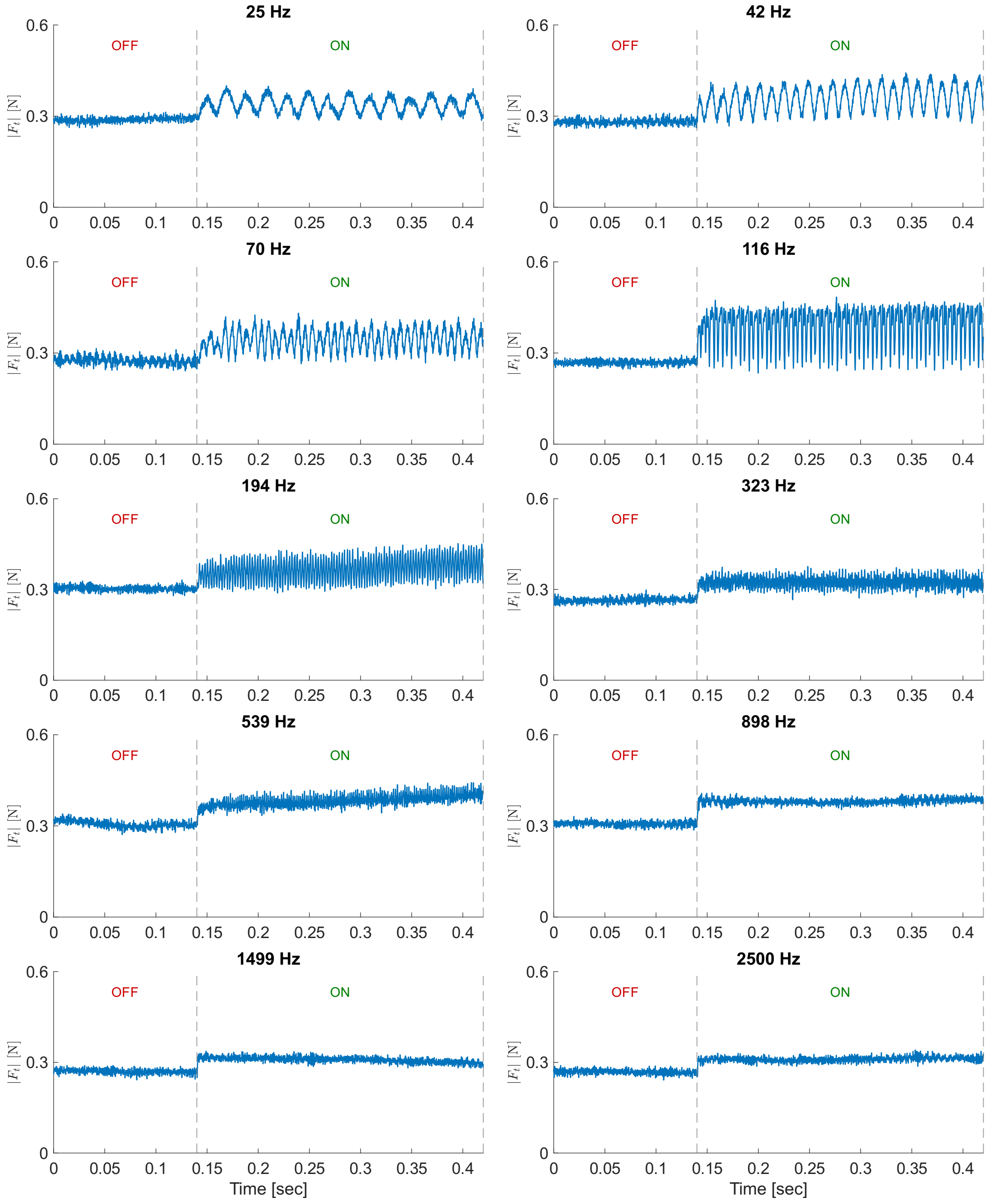


**Figure S15: Tangential force measurements for all frequencies.** Representative tangential force measurements for all frequencies when the voltage is off and on from Participant-1.

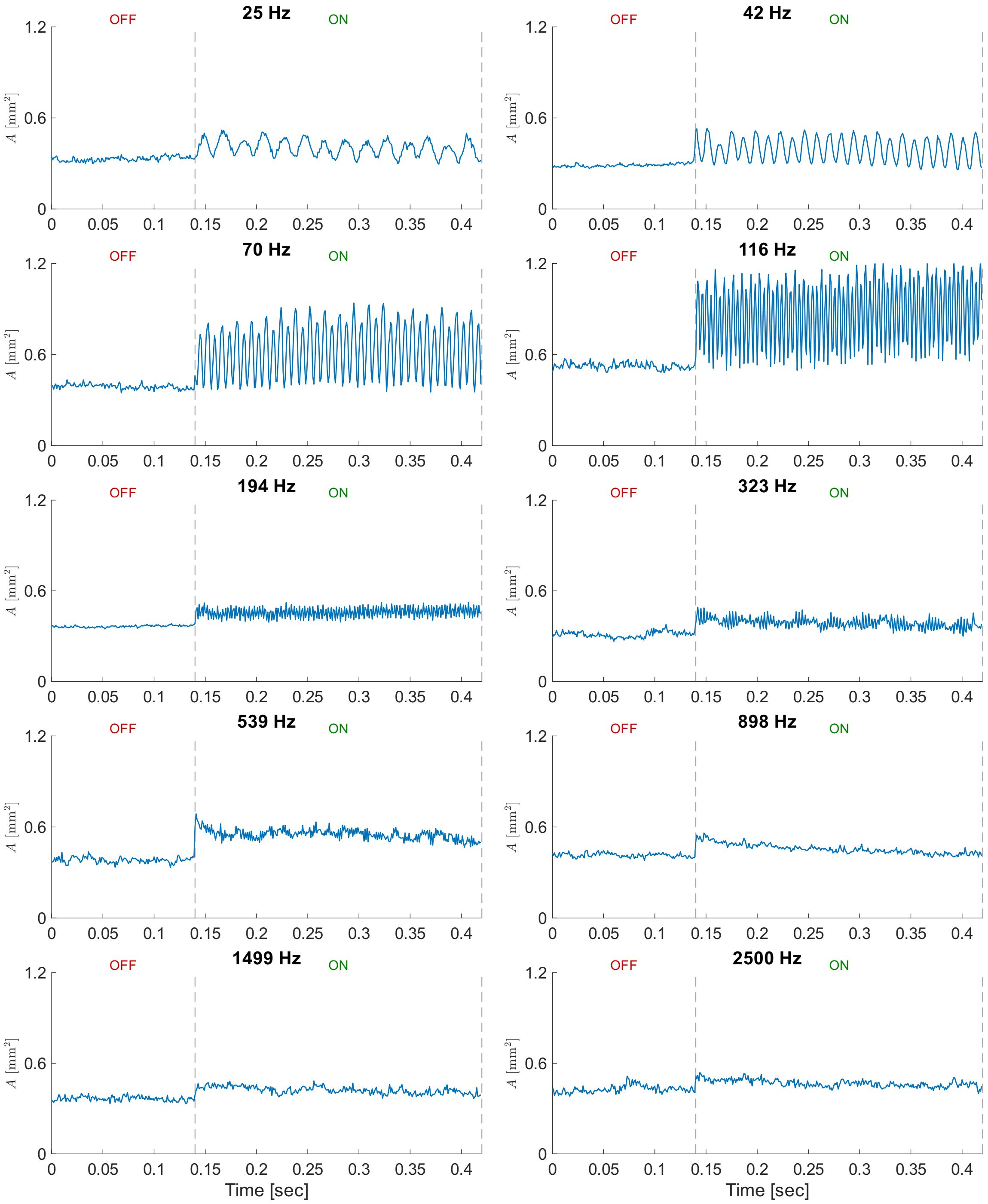


**Figure S16: Real area measurements for all frequencies.** Representative real area measurements for all frequencies when the voltage is off and on from Participant-1.

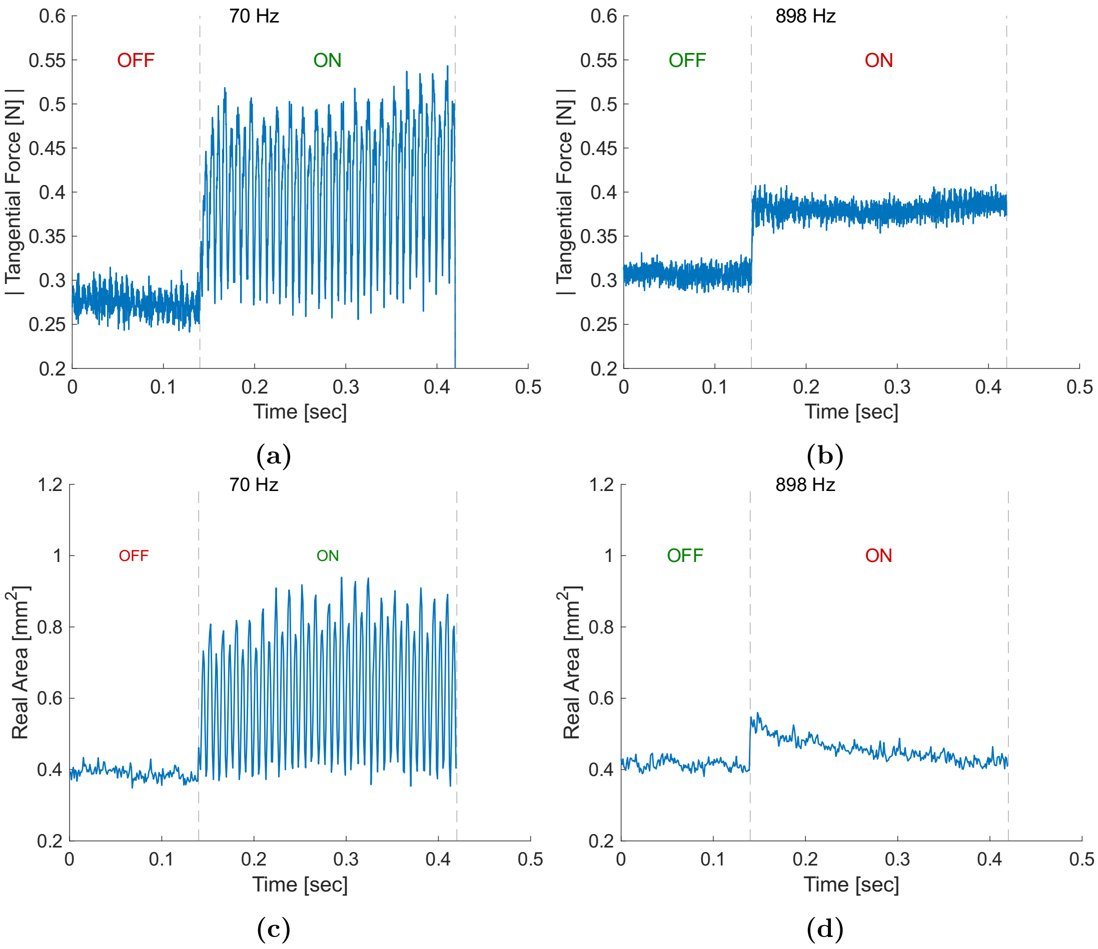


**Figure S17: Example measured tangential force and real area from vibration and adhesion regime.** Example tangential force data recorded during a trial when (a) $f_0 = 70$ Hz and (b) $f_0 = 898$ Hz. Example real area data recorded during a trial when (c) $f_0 = 70$ Hz and (d) $f_0 = 898$ Hz. Oscillation is observed for both signals in the vibration regime, but not in the adhesion regime.

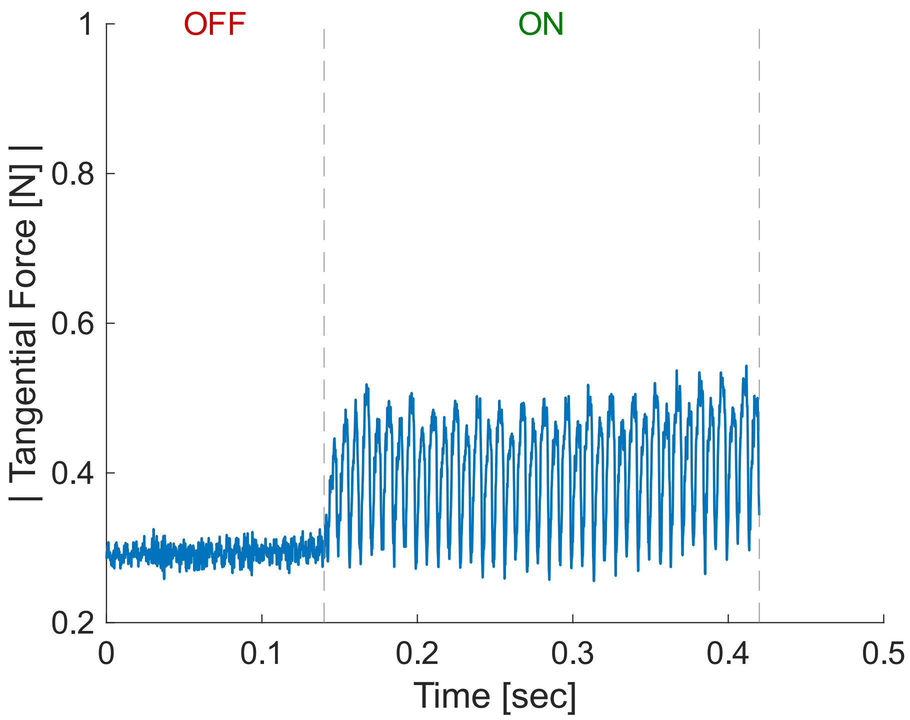


**(a)**

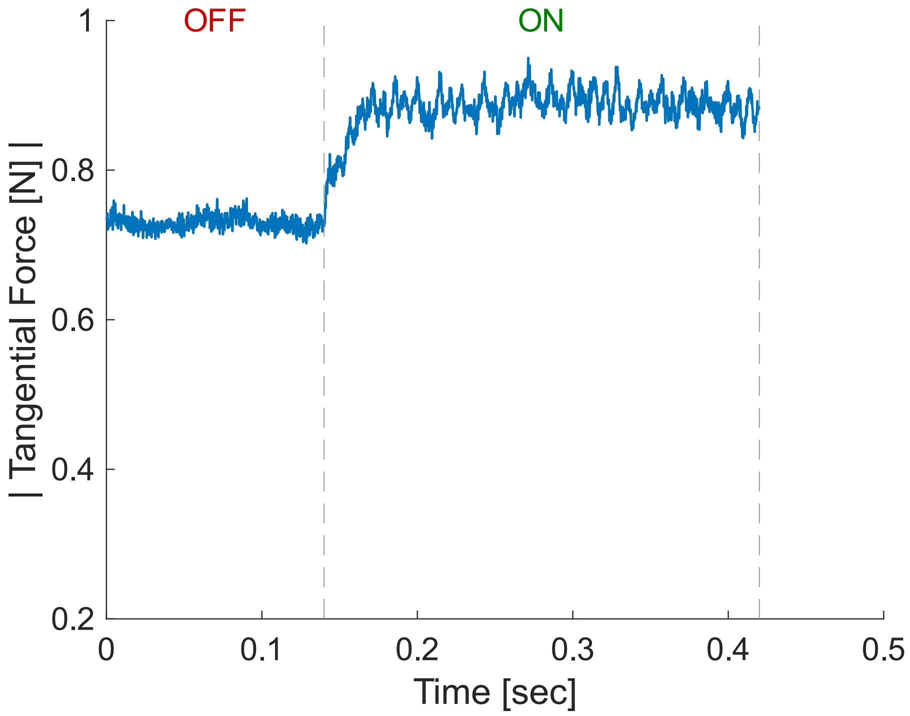


**(b)**

**Figure S18: Comparison of low-frequency oscillatory responses between two participants.** (a-b) Tangential force modulation of 2 participants in vibration regime ($f_0 = 70$ Hz). One participant demonstrates a distinct oscillatory modulation, while another exhibits a considerably weaker modulation.

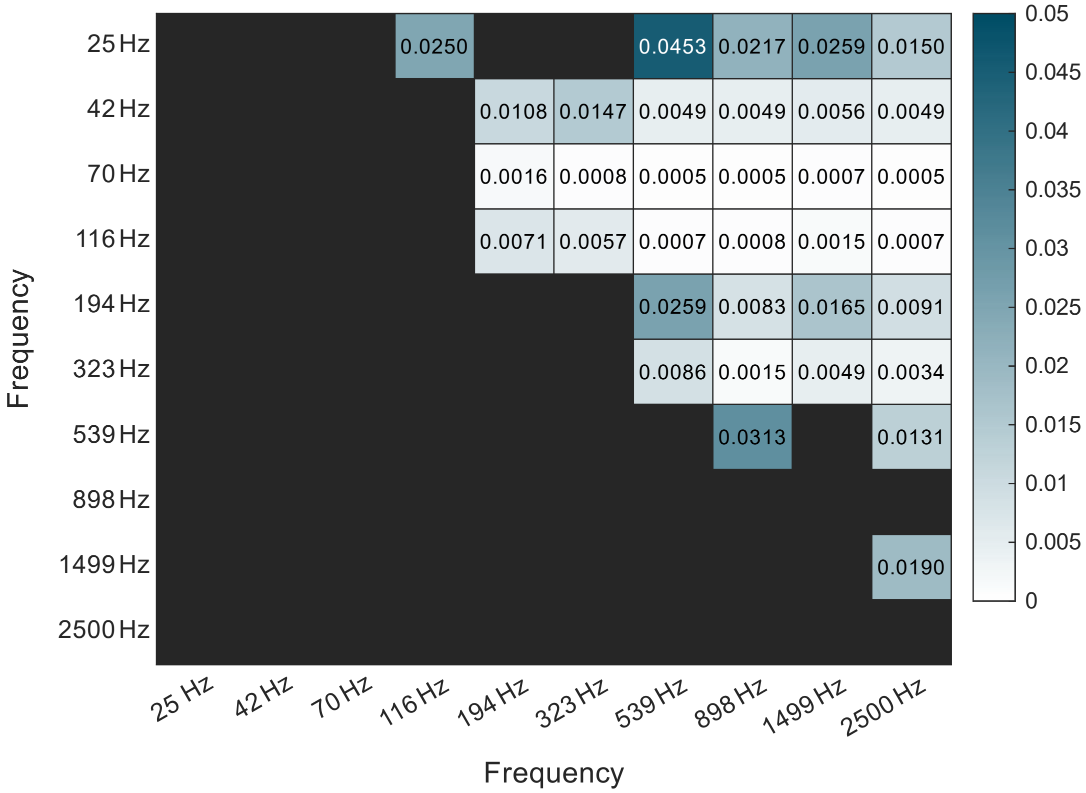


**Figure S19: Post hoc paired t-tests for interfacial shear stress ratio.** Results of the post hoc paired t-tests with Benjamini-Hochberg for interfacial shear stress ratios ($\tau^{on}/\tau^{off}$) across frequencies

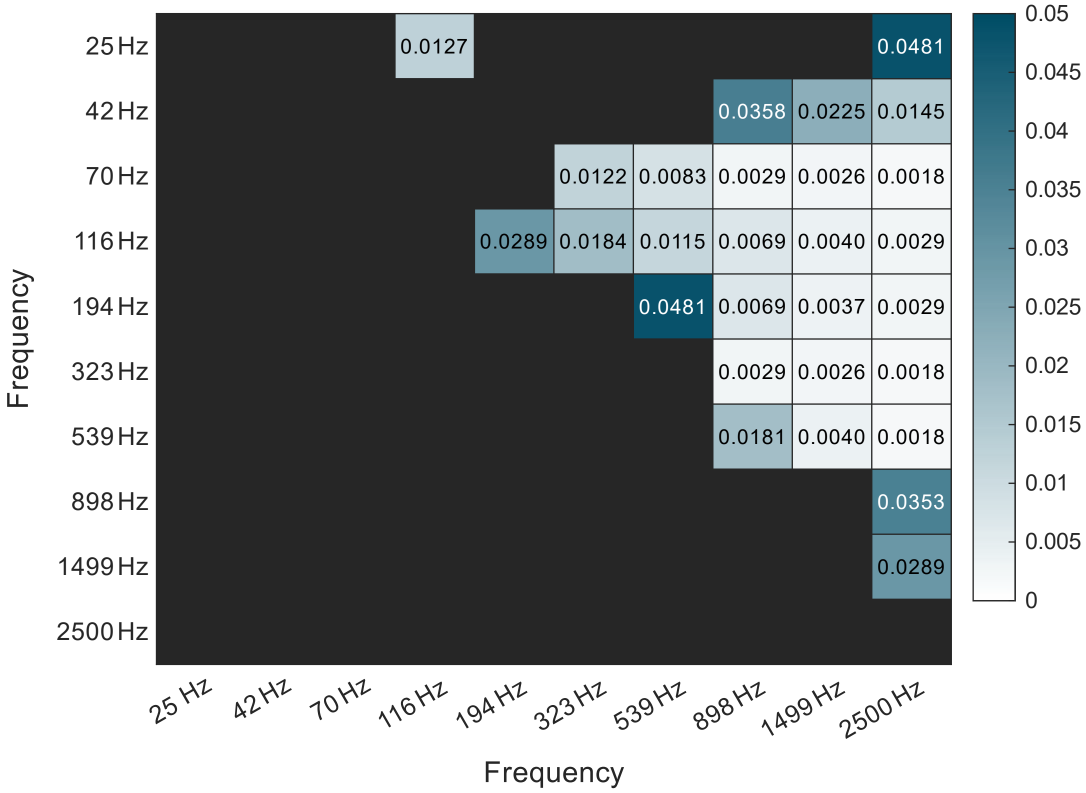


**Figure S20: Post hoc paired t-tests for real area ratio.** Results of the post hoc paired t-tests with Benjamini-Hochberg for real area ratios $(A^{on}/A^{off})$ across frequencies

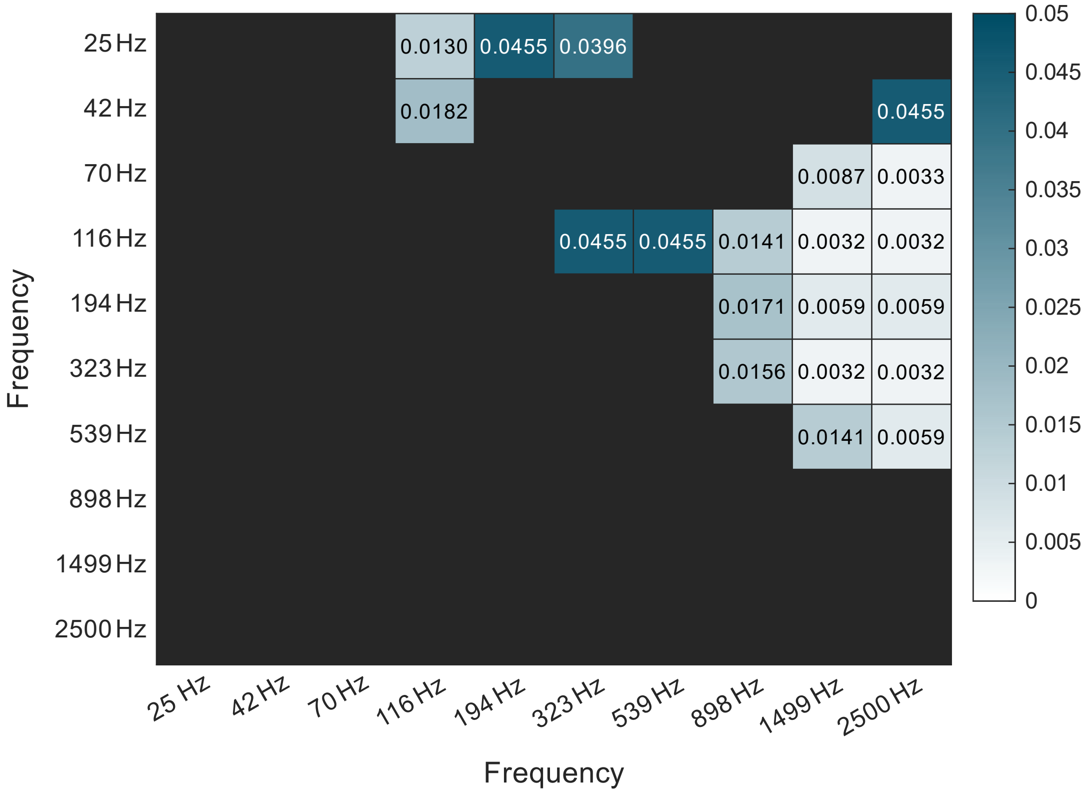


**Figure S21: Post hoc paired t-tests for tangential force ratio.** Results of the post hoc paired t-tests with Benjamini-Hochberg for tangential force ratios ($F_t^{on}/F_t^{off}$) across frequencies.

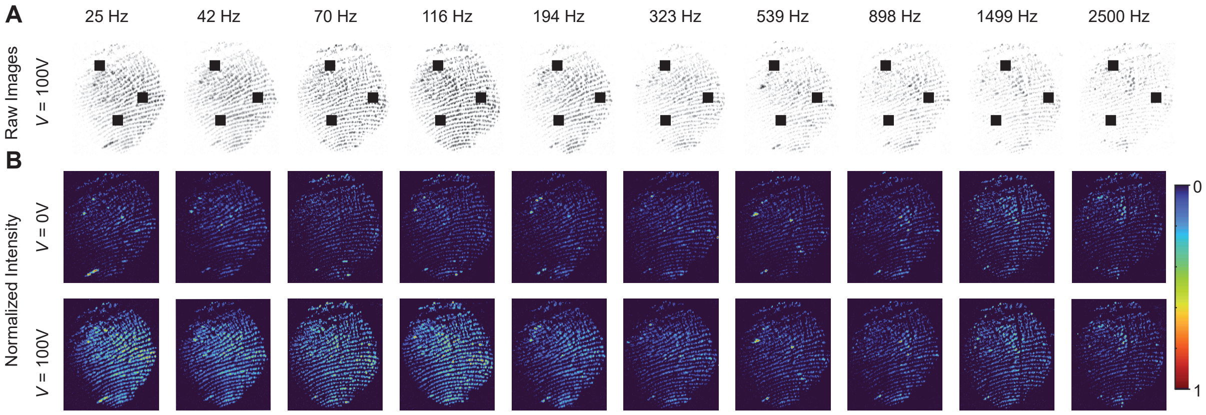


**Figure S22: Example fingertip contact images across frequencies.** (A) Raw contact area images from a single participant under 100 V electrostatic actuation at various frequencies. Darker regions correspond to greater real contact area. (B) Normalized contact intensity maps comparing 0 V and 100 V conditions. For visualization, pixel intensities were normalized within each frequency by the maximum pixel value observed.

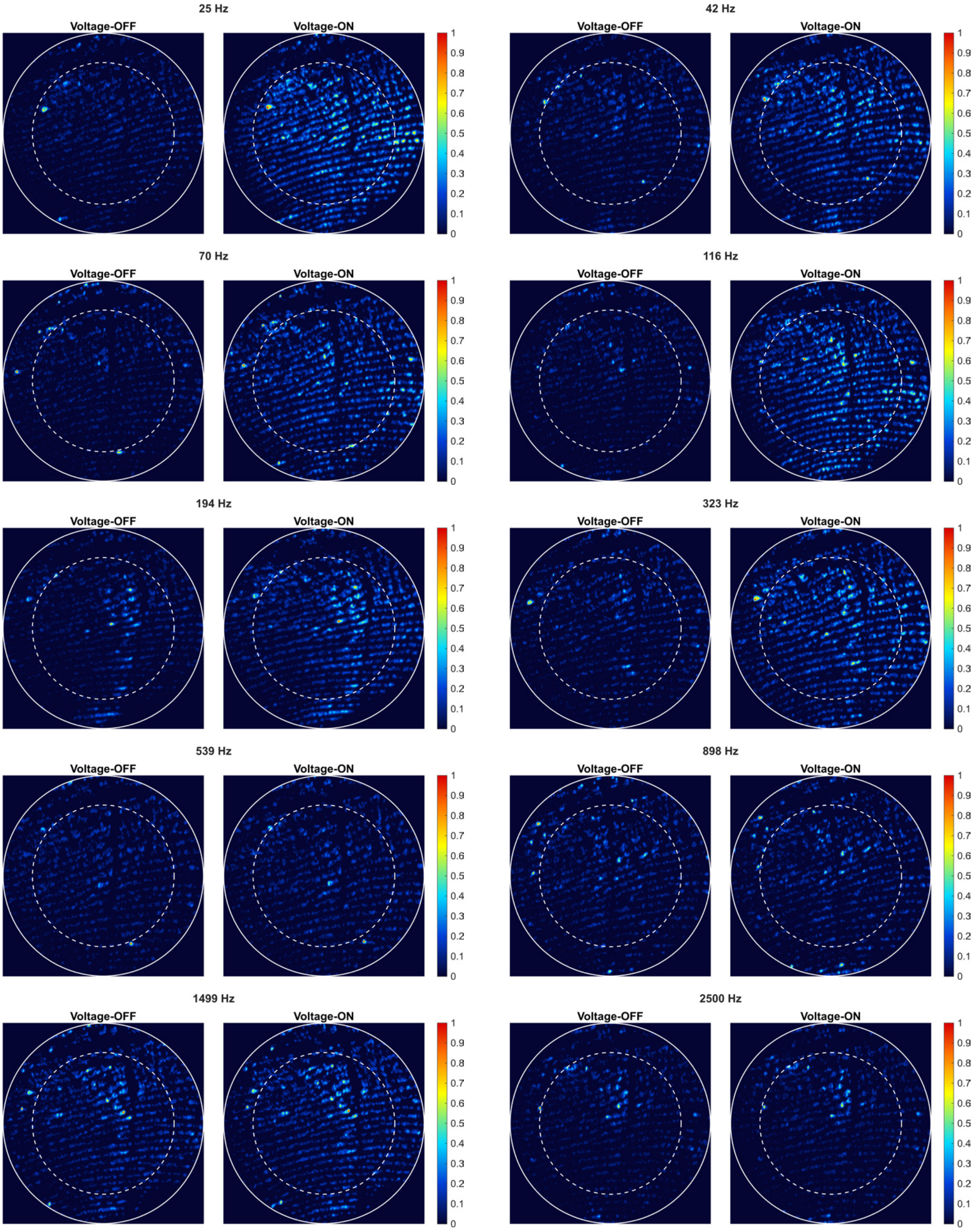


**Figure S23: Example normalized spatial contact maps of the fingertip contact region for the voltage-off ($V = 0$) and voltage-on ($V = 100$) conditions across all tested frequencies.** For each frequency, the corresponding voltage-off ($V = 0$) and voltage-on ($V = 100$) maps were normalized together using a common intensity scale, allowing direct comparison between the two conditions within that frequency. The dashed contour indicates the boundary between the central and peripheral regions used for the qualitative spatial comparison.

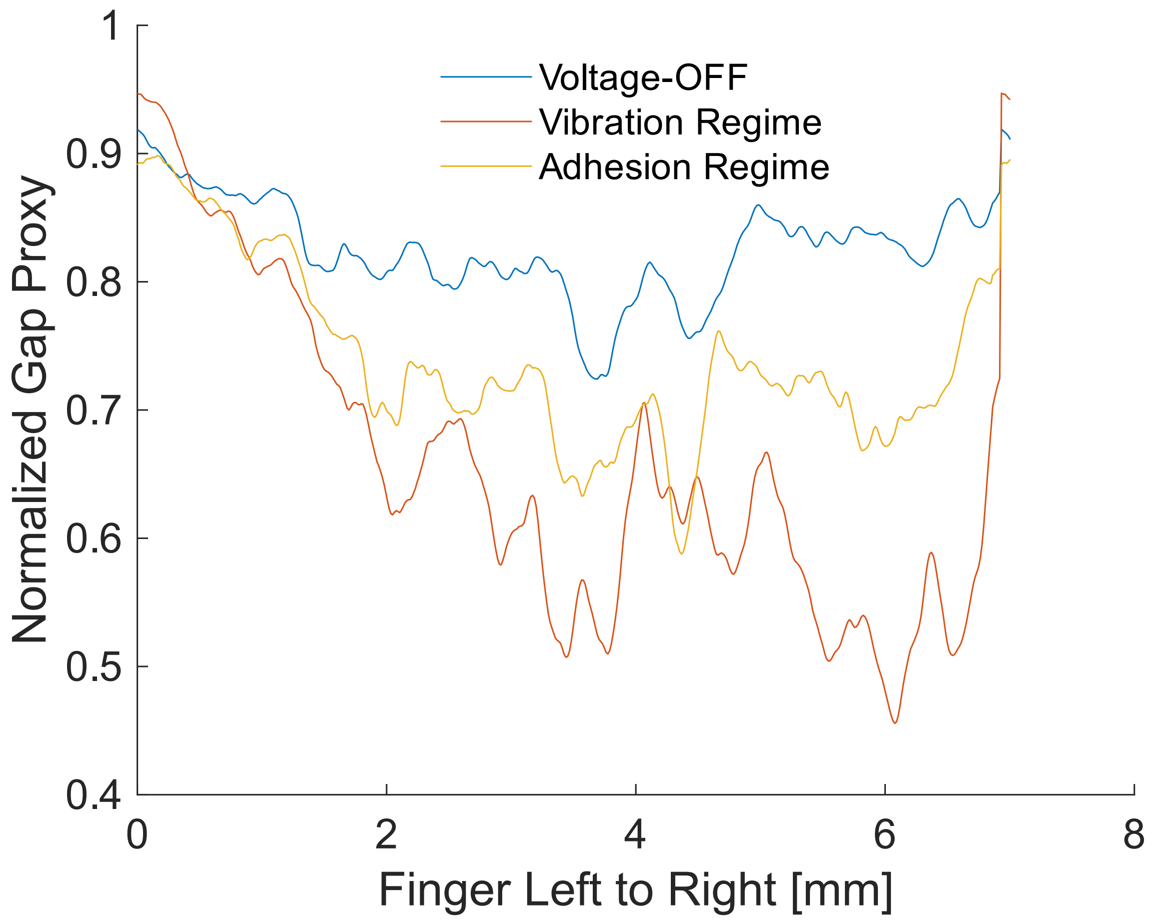


**Figure S24: Relative gap profiles across the finger contact.** Relative interfacial gap profiles extracted from the middle of the fingerprint images under voltage-off ($V = 0$) and voltage-on ($V = 100$) conditions. The profiles were obtained by averaging pixel values column-wise within a 1 mm-high horizontal rectangular region placed across the middle of the fingerprint image. Profiles were normalized within each frequency and then averaged within the vibration and adhesion regimes. Lower values indicate a smaller relative gap.

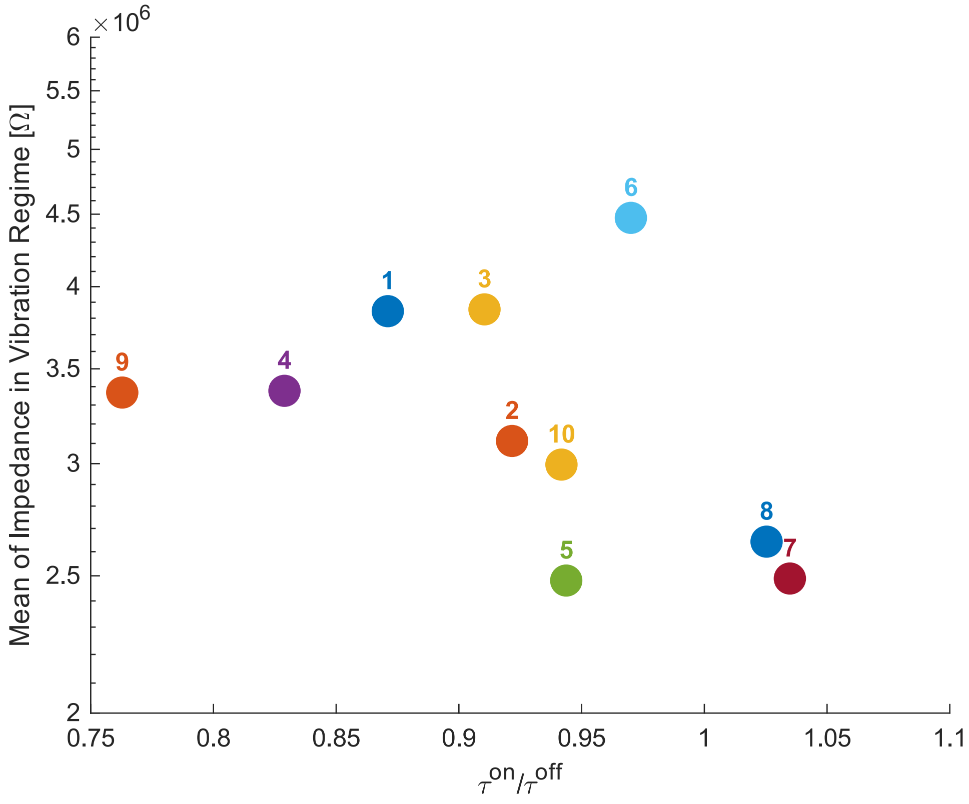


**Figure S25: Measured electrical impedance in the vibration regime.** Mean electrical impedance within the vibration regime for each participant.

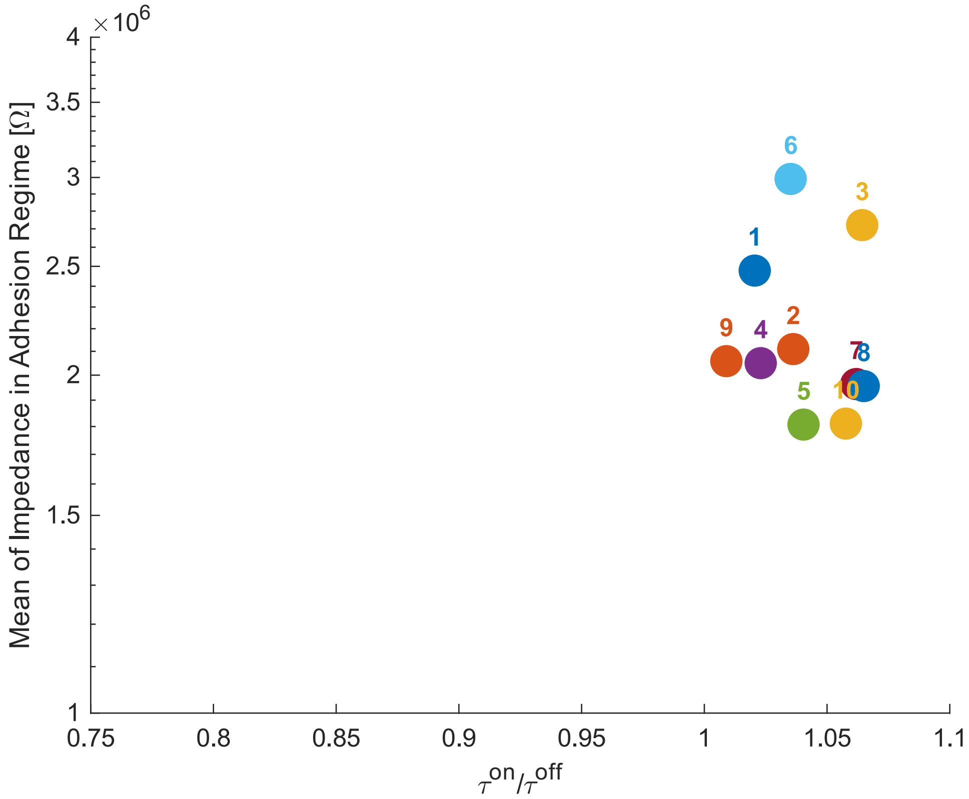


**Figure S26: Measured electrical impedance in the adhesion regime.** Mean electrical impedance within the adhesion regime for each participant.

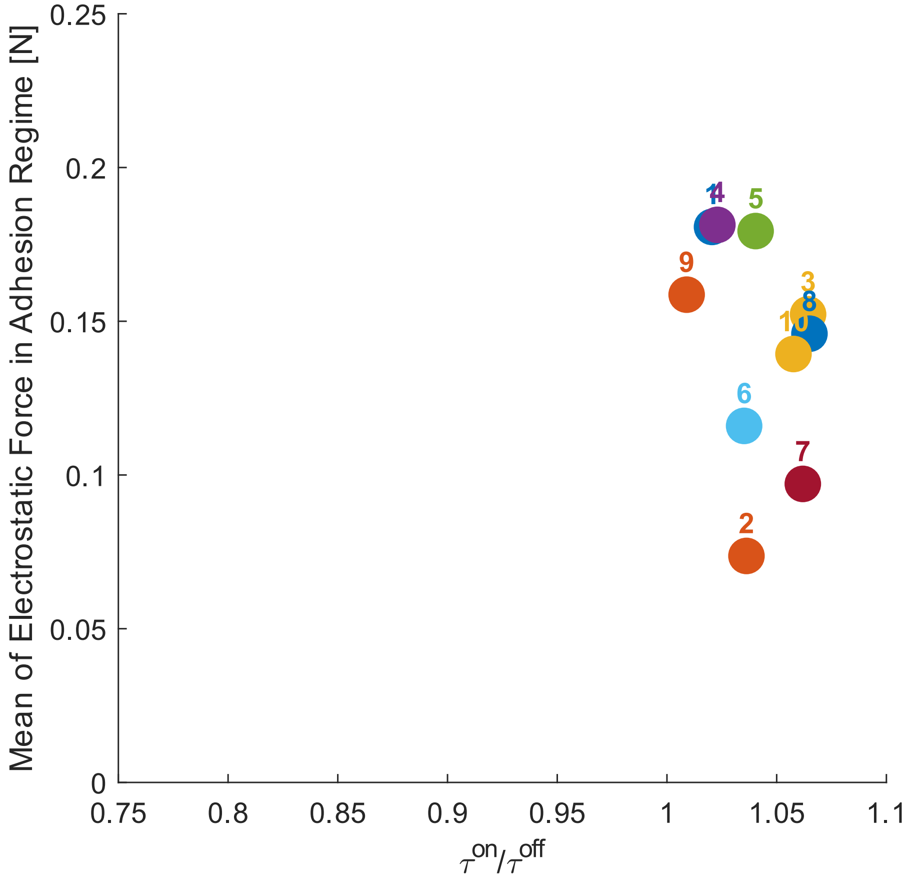


**Figure S27: Measured electrostatic force in the adhesion regime.** Mean electrostatic force within the adhesion regime for each participant.

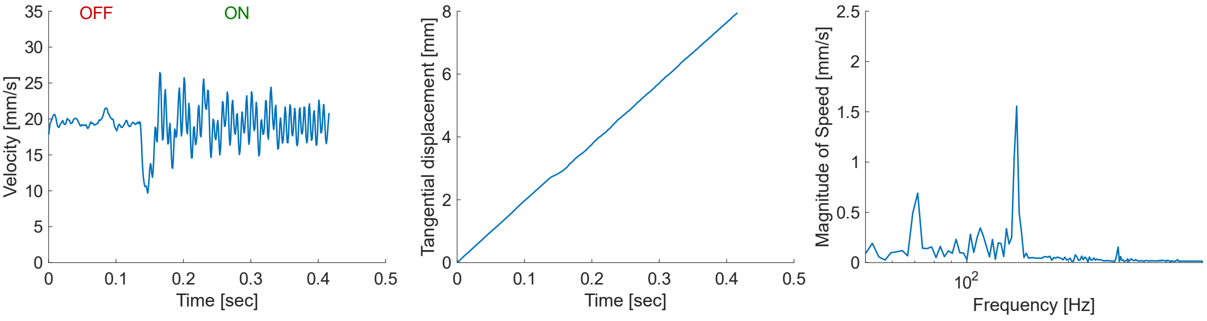


**(a)**

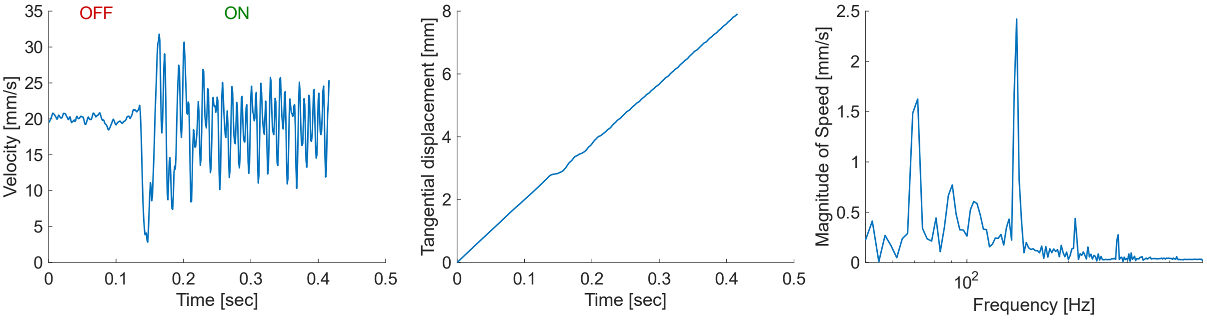


**(b)**

**Figure S28: Example analysis of tangential finger motion from supplementary videos.** (a) Extracted tangential velocity, displacement, and velocity spectrum from a participant in Movie S1. (b) Extracted tangential velocity, displacement, and velocity spectrum from a participant in Movie S2. Instantaneous tangential velocity and displacement traces extracted from the supplementary videos at low actuation frequency ($f_o = 70$ Hz), together with their corresponding velocity spectra. The motion shows clear within-cycle velocity modulation, with a pronounced spectral component at the effective actuation frequency ($2f_0$), consistent with partial stick-slip-like tangential modulation.

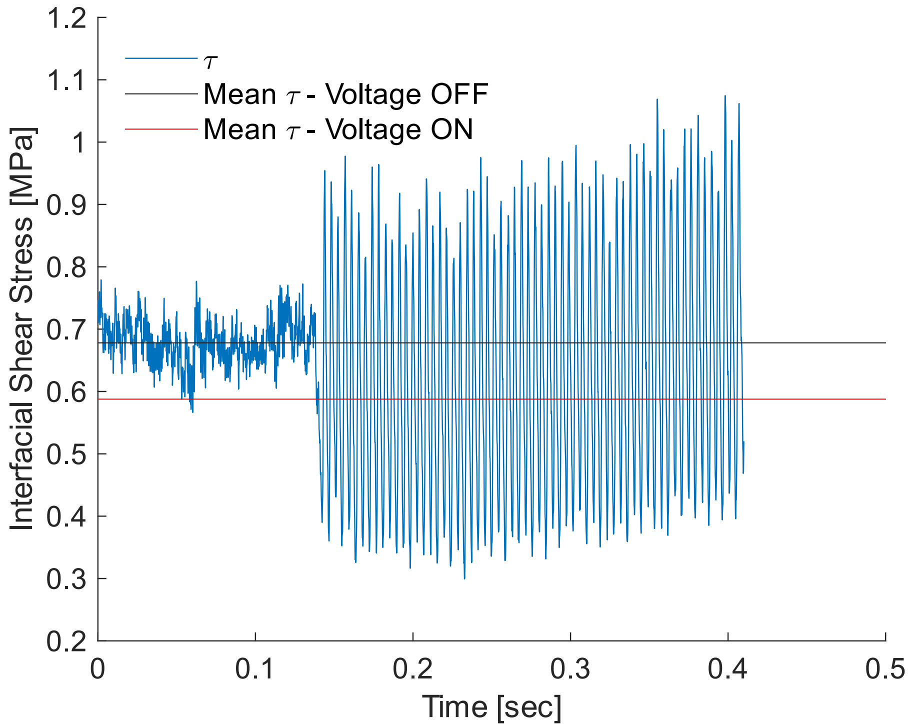


**Figure S29: Representative time-resolved interfacial shear-stress.** Representative time-resolved interfacial shear-stress estimate for one participant, calculated from the measured tangential force and optically resolved real contact area.

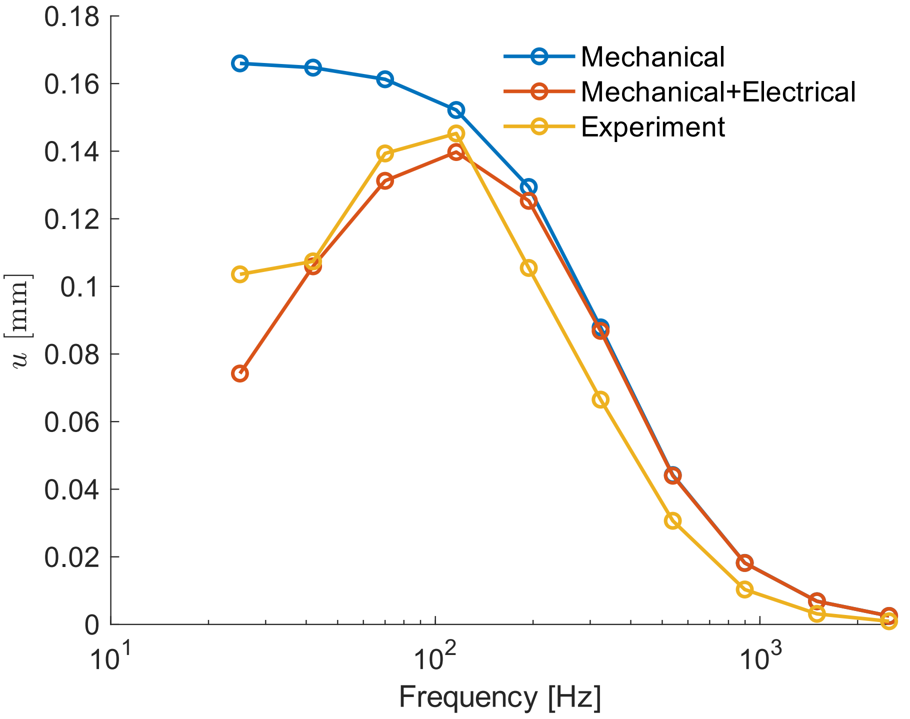


**Figure S30: Effect of the electrostatic forcing term on the predicted peak frequency.** Comparison of the spring–mass–damper model using three electrostatic force inputs: constant amplitude, amplitude weighted to reflect a first-order electrical contribution, and experimentally inferred $F_e$. The predicted peak frequency depends on the assumed force input, suggesting that the observed 116 Hz peak reflects coupled electrical–mechanical behavior rather than a purely mechanical resonance.

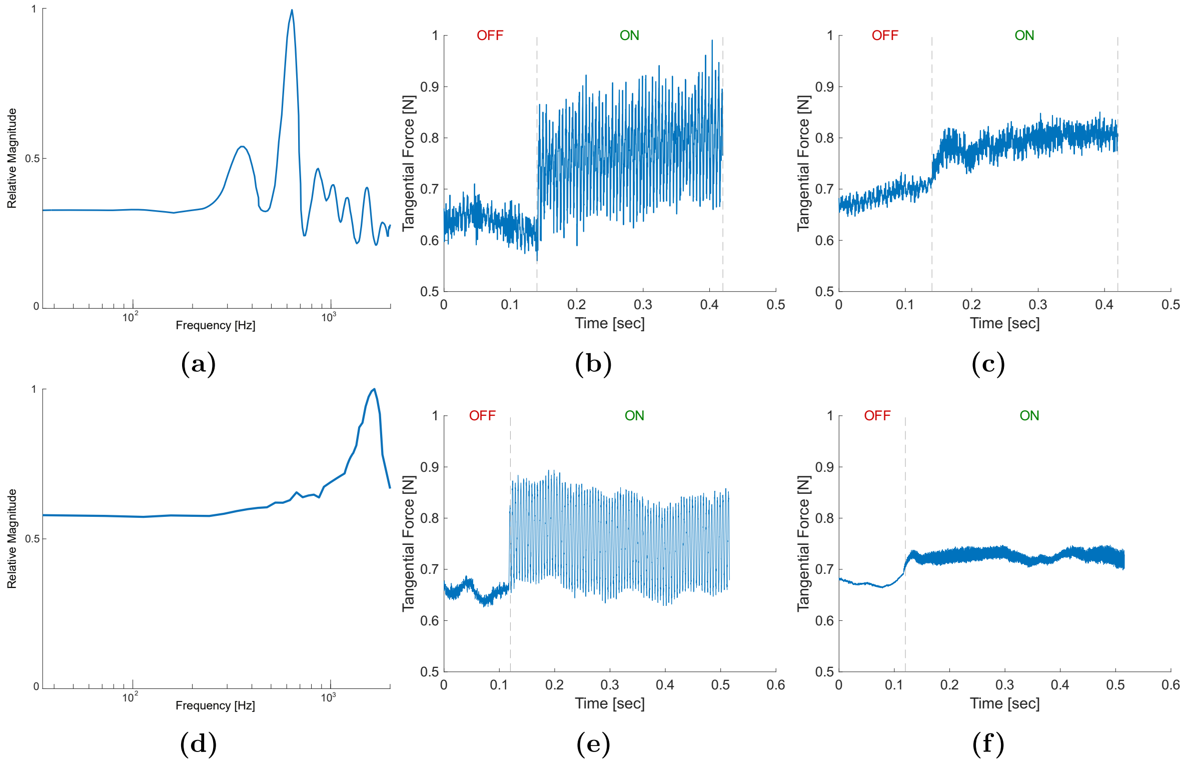


**Figure S31: Comparison between the dynamic characterization of the experimental setup and its corresponding tangential force responses, and those of a reference (verification) setup.** (a) Impact-hammer test result of the experimental setup in the tangential direction. (b,c) Representative tangential-force traces from the experimental setup at $f_0 = 100$ Hz and $f_0 = 600$ Hz. (d) Impact-hammer test result of the verification setup in the tangential direction. (e,f) Representative tangential force traces from the verification setup at $f_0 = 100$ Hz and $f_0 = 600$ Hz.

#### Supplementary Movies

**MovieS1** Video of the fingerprint contact during one trial when the input voltage was at 70 Hz (example 1)

**MovieS2** Video of the fingerprint contact during one trial when the input voltage was at 70 Hz (example 2).

**MovieS3** Video of the fingerprint contact during one trial when the input voltage was at 539 Hz (example 1).

**MovieS4** Video of the fingerprint contact during one trial when the input voltage was at 539 Hz (example 2).

**MovieS5** Video of the moist fingerprint contact during one trial when the input voltage was at 42 Hz.
